# Towards a fine-scale population health monitoring system

**DOI:** 10.1101/780668

**Authors:** Gillian M Belbin, Stephane Wenric, Sinead Cullina, Benjamin S Glicksberg, Arden Moscati, Genevieve L Wojcik, Ruhollah Shemirani, Noam D Beckmann, Ariella Cohain, Elena P Sorokin, Danny S Park, Jose-Luis Ambite, Steve Ellis, Adam Auton, CBIPM Genomics Team, CBIPM Genomics Team, Regeneron Genetics Center, Erwin P. Bottinger, Judy H Cho, Ruth JF Loos, Noura S Abul-husn, Noah A Zaitlen, Christopher R Gignoux, Eimear E Kenny

## Abstract

Understanding population health disparities is an essential component of equitable precision health efforts. Epidemiology research often relies on definitions of race and ethnicity, but these population labels may not adequately capture disease burdens specific to sub-populations. Here we propose a framework for repurposing data from Electronic Health Records (EHRs) in concert with genomic data to explore enrichment of disease within sub-populations. Using data from a diverse biobank in New York City, we genetically identified 17 sub-populations, and noted the presence of genetic founder effects in 7. By then linking community membership to the EHR, we were able to identify over 600 health outcomes that were statistically enriched within a specific population, with many representing known associations, and many others being novel. This work reinforces the utility of linking genomic data to EHRs, and provides a framework towards fine-scale monitoring of population health.

## Introduction

Populations around the world often have differential rates of disease due to a combination of genetic variation and environmental exposures. Understanding the differences in disease burdens according to demographic factors is the basis of epidemiology research, and is fundamentally important to clinical and population health practice. Most studies of human disease begin by sampling from predefined populations, which are usually identified on the basis of race, ethnicity, cultural identity or geography. However, the choice of population categories may incompletely capture the environmental and demographic ties that can impact disease burdens. In the United States, individuals with roots from Latin American are often classified as Hispanic and/or Latino, but sub-groups with origins from different countries in the Americas may have different rates of disease. For example, populations of Puerto Rican descent have one of the highest asthma rates in the world, while populations of Mexican descent have one of the lowest^1,2^.

With the advent of large, population-based DNA biobanks in health systems, new opportunities are available to characterize the links between demography and a broad range of health outcomes^3–7^. Knowledge about genetic variation shared across human populations can aid in understanding the demographic events that might impact disease burden across populations. For example, risk variants in the *APOL1* gene, which confer a substantially increased risk of kidney disease and cardiovascular disease, arose in Africa, were first discovered in African American populations^8,9^, and are mainly studied in African or African American populations. However, *APOL1* risk variants exist at appreciable frequencies among many populations across the Americas who historically share African genetic ancestry, but do not self-identify as African or African American, and are subsequently underrepresented in *APOL1* research^10,11^. This suggests, that while self-reported population information is useful in assessing epidemiological risk, in some cases it may be limiting. Furthermore, self-reported population demographic information may be encoded inaccurately in health systems, can be limited in scope, and therefore may not accurately recapitulate the inherent population structure impacting disease risk^12,13^.

High density, genome-scale data has long been used to examine genetic differences between populations, which in turn, can be used to infer population genetic history. Popular techniques are algorithmic-based methods such as principal component analysis (PCA)^14,15,16^, or model-based methods such as ADMIXTURE^17–19^, which estimate genetic ancestry by assessing genomic variants in aggregate. Other powerful approaches infer more fine-scale genetic ancestry by using local haplotypes along chromosomal segments^20–22^. One such approach identifies residual signatures of distant genealogical relationships which may be detected by inferring long range haplotypes^23^. Although genetic traces of our distant ancestors will decay rapidly over time, long haplotypes that have been co-inherited Identical-by-Descent (IBD) from some of our more recent ancestors may persist. At a population level these can be analyzed in aggregate to infer distant relationships to a set of shared ancestors. We expect that individuals in a population who are genealogically closer to one another may also be more likely to share recent population history. This in turn may be linked to sharing of culture, environment and correlations in disease risk. Furthermore, elevations in IBD sharing within population groups is also indicative of recent founder effects which can impact prevalence of disease variants. Therefore, examining fine-scale population structure as inferred using genetic genealogical data, in particular when linked to electronic health records (EHRs) in health systems, offers new opportunities to explore genome-phenome architecture and health disparities.

In this study, we examined fine-scale population structure in Bio*Me*, a highly diverse, multi-ethnic biobank ascertained through the Mount Sinai health system in New York City (NYC). We explored how demographic information is currently captured in the EHR in the form of recorded ethnicity. We compared EHR-recorded ethnicity with self-reporting information volunteered by over 36,000 Bio*Me* participants. We discovered high levels of discordance between EHR-recorded and self-reported ethnicity for some populations in NYC. Exploring genetic ancestry within Bio*Me* revealed further complexity, with distinct patterns of continental and subcontinental genetic substructure within self-reported ethnic groups. To explore the substructure further, we analyzed IBD sharing in Bio*Me* and applied an unsupervised, scale-free network modeling method to uncover clusters of populations informed by patterns of recent demography. We reveal 17 distinct communities enriched for recent, shared genealogy and highly correlated with recent migratory patterns to NYC. We demonstrated that many of these community’s harbor signatures of founder events, the timings of which coincided with the era of colonization of the Americas. Finally, we linked these communities to over 1700 health outcomes and demonstrate distinctive health patterns of disease risk, uncovering over 2000 examples of statistically significant differences in health outcomes between populations, some of which point to unknown or underappreciated population-specific disease risks. This work demonstrates the utility of the application of genomic data towards fine-scale population health monitoring.

## Results

### Evaluating ethnicity as captured in the large Mount Sinai health system in New York City

We evaluated ethnicity in a large (N=36,061), diverse biobank (Bio*Me*) linked to electronic health records (EHRs) in the Mount Sinai Health System in NYC. To understand how ethnicity information is captured in a large, urban health system, we compared the self-reported ethnicity surveyed during the Bio*Me* enrollment (self-reported ethnicity; **Supplementary Table 1**) to the ethnicity recorded in the Mount Sinai EHR. We first restricted analysis to participants who reported only one ethnicity in the enrollment survey for one of five population groups: European American (EA, N=9830), African-American (AA; N=7976), Hispanic/Latino (HL; N=11544), East and Southeast Asian (ESA; N=965), and Native American (NA; N=61), (total: 84.2% of BioMe; N=30376). We mapped self-reported ethnicity for these individuals to ethnicity information extracted from the electronic health records at each independent visit (n=1310279 total visits). Each participant had visited the health system between years 2007-2014 between a median of 22 (range 1-1076) times (**Supplementary Figure 1**).

Self-reported and EHR-recorded ethnicity were concordant 64.5% of total visits, or 71.6% of visits if excluding visits where ethnicity was designated as ‘Unknown’ (resulting in the exclusion of n=129678 healthcare visits). We observed significant differences in ethnicity concordance between population groups (**Figure 1A**), with significant differences (ANOVA 2< p×10-16) in concordance between ethnicities; EA (median=91.2%; 95% confidence intervals=90.2%-92.0%), AA (100%; 100%-100%), HL (53.7%; 52.9%-54.4%), ESA (63.6%; 55.6%-66.7%), and NA (0%,0%-0%). However, only 29.8% of patients were concordant with self-reported ethnicity in all their visits, and we observed differences between population groups; 34.4% EA, 51.4% AA, 12% HL,21.7% ESA, and 5% NA. This concordance increases when we require that patients be concordant in only half of their visits, however, there remain notable difference between groups; 91% AA, 78% EA, 58.2% ESA and 56.2% HL, and 13.3% NA (**Figure 1B)**.

**Figure 1.**
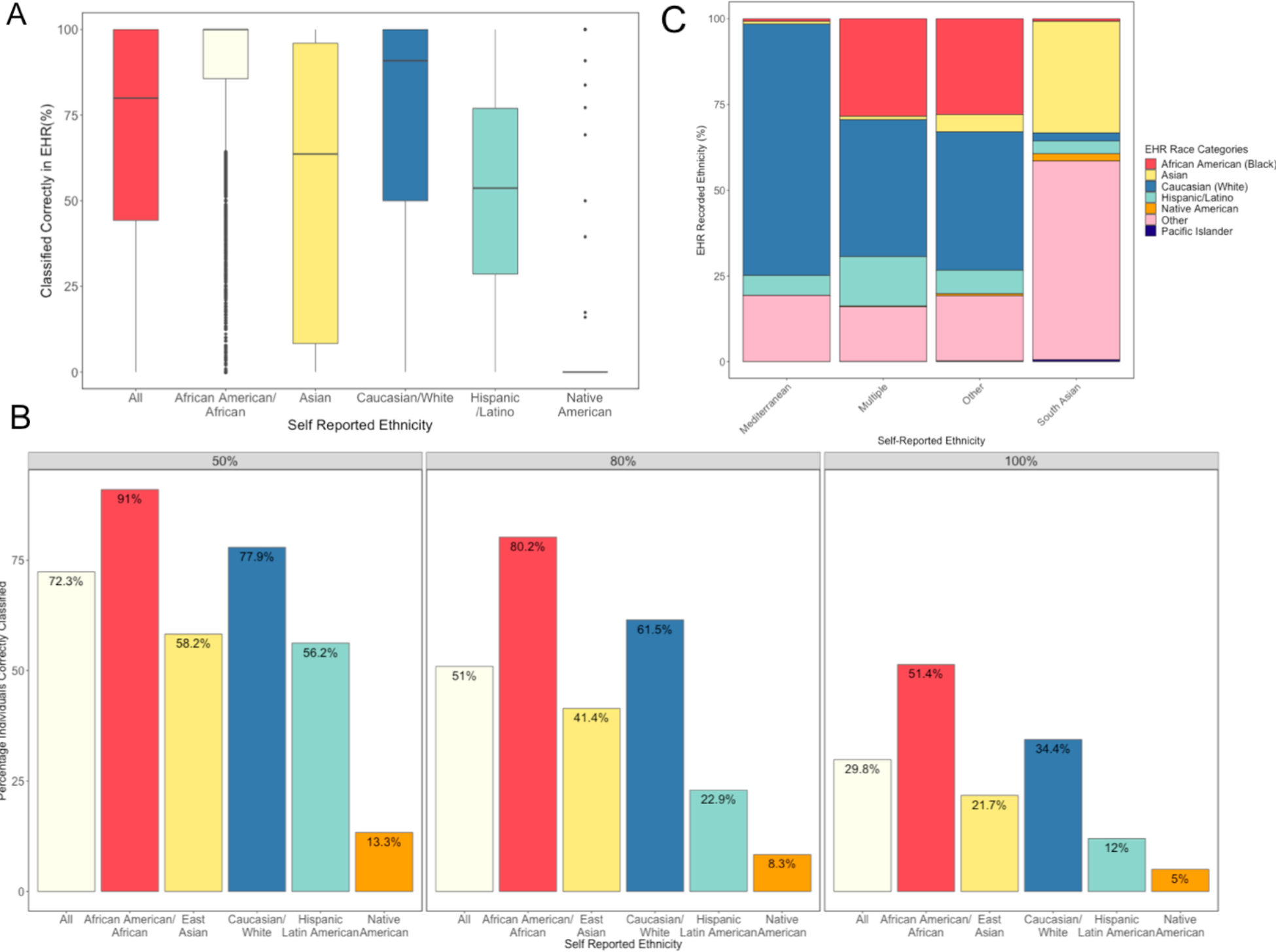
(**A**) Distribution of the percentage of times an individual’s self-reported ethnicity matches their EHR designated ethnicity, across all individuals and stratified by self-reported ethnicity. (**B**) Proportion of individuals correctly classified across 50%, 80% and 100% of visits. (**C**) EHR ethnicity designations for individuals whose self-reported ethnicity did not correspond to one of the EHR categories.

15.8% of participants self-reported a different, namely South Asian, Mediterranean, Other, or Multiple ethnicities at enrollment (N=5685 individuals across 182149 healthcare visits). Individuals who self-reported as South Asian were most often classified as “Other” (45.9%) or “Asian” (25.8%). Individuals who self-identified as “Mediterranean” were most often classified as “Caucasian” (59.6%) or “Other” (15.7%), while for individuals who either self-reported as “Other” or who checked multiple categories there was no clear majority designation (**Figure 1C**). Overall these analyses support that ethnicity data is often poorly and inconsistently captured, particularly for non-EA populations, during hospital visits in the Mount Sinai health system.

### Characterizing genetic diversity in the BioMe Biobank

To explore the relationship between self-reported ethnicity and genetic ancestry we estimated global genetic ancestry proportions for a subset of Bio*Me* participants (N=31705) genotyped on the Global Screening Array (GSA). We merged these individuals with a reference panels of 87 populations representing ancestry from 7 continental or subcontinental regions. Using PCA, we demonstrate that Bio*Me* participants represent a continuum of genetic diversity between African and non-African reference panels on principal component (PC) 1, and between European, Asian, American and Oceanian reference panels on PC 2 **(Figure 2A**). The first 10 PCs can also be represented as a low-dimensional topological map using the Uniform Manifold Approximation and Projection (UMAP) algorithm, presenting many clusters that are roughly grouped by continent and others with little or no overlap with reference panels presenting a challenge for their interpretation (**Figure 2B)**.

**Figure 2.**
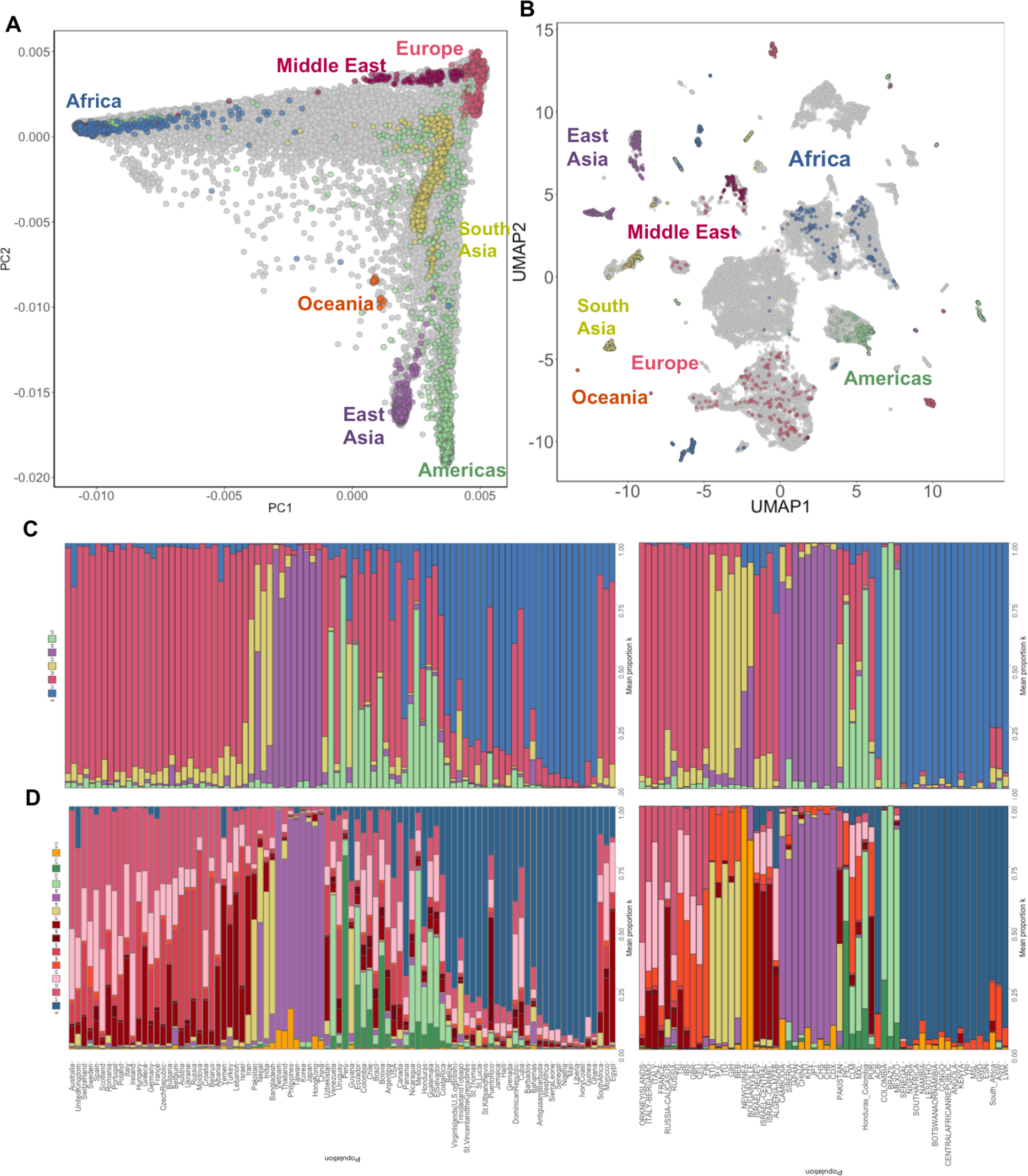
(**A**) Principal Component Analysis (PCA) of N=31705 BioMe participants (grey) along with N=4149 reference panel individuals representing 7 Continental/Subcontinental regions, namely Africa (blue), Europe (red), The Middle East (dark red), South Asia (yellow), Oceania (orange), East Asia (purple) and the Americas (green). Bio*Me* participants can be observed to fall across the spectrum of global diversity, with many participants falling on a cline between various Subcontinental reference panels, suggesting the presence of varying degrees of admixture. (**B**) Projection of UMAP across Bio*Me* (grey) and continental reference panels based on PC1-PC10 reveals patterns of fine-scale genetic substructure. (**C**) Stacked bar chart representing mean admixture proportions per country for Bio*Me* participants (N>= 10) (left) and reference panels (right) obtained by running ADMIXTURE at *k=5* reveals varying proportions of European (red), South Asian (yellow), East Asian (purple), African (blue) and Native American (green) in Bio*Me*. (**D**) Running ADMIXTURE at *k=12* reveals fine-scale patterns of subcontinental ancestry, including an Ashkenazi Jewish component (dark red), and a drifted Iberian component (light pink) that is particularly notable in Puerto Rican born Bio*Me* participants.

We also explored population demography using a model-based approach called ADMIXTURE that applies a pre-set number of putative ancestral populations to seek the best fit of ancestral clusters in the data (**Supplementary Figure 2**). Analysis that fit five ancestral populations (*k=5*) recapitulates five continental level components corresponding to African, European, East Asian, South Asian and Native American ancestry. As expected, self-reported AA and H/L participants exhibited varying degrees of ancestry from Europe, Africa and the Americas, however, several individuals also appear to have appreciable (>10%) levels of South Asian (2.2% of individuals) or East Asian (0.6% of individuals) ancestry. Participants who self-identify as ‘Other’ exhibit varying proportions of admixture from all 5 different continent groups (**Supplementary Table 2**).

By limiting the analysis to Bio*Me* participants born outside continental US (35% of participants), we were able to explore diversity linked to more recent demography (**Figure 2C**). We demonstrate European, African and Native American ancestral proportions consistent with previous reports in participants born in Puerto Rico (59.3%,26.0%,14.2% respectively), Dominican Republic (50.3%, 40.9%, 7.4%), Jamaica (13.0%, 83.2%, 0.6%) and Cuba (66.4%, 26.8%, 4.6%)^24^. We observe appreciable levels of East and South Asian ancestry in participants born in Trinidad and Tobago (4.7%,24.5%), the Bahamas (10.3%,7.7%) and Guyana (6.1%,50.6%), which is consistent with historical accounts of South Asian migration to the Caribbean. At *k=12* **(Figure 2D)** we also observe complex structure in the European component including the resolution of an Ashkenazi Jewish component in a subset of the US-born participants, and two distinct European components that likely represent a Northern-Southern European genetic cline, and a distinct European component that appears predominantly in admixed populations from the Americas. This component is particularly notable in Caribbean-born participants and is present to a lesser extent in Portuguese and Spanish born Bio*Me* participants. Taken together, this information reveals Bio*Me* to be a rich source of ancestral genetic diversity, including many populations that are otherwise poorly represented in biomedical genomics research.

### Detecting Communities of Recent Shared Ancestry in New York City

To investigate the demographics of migration to New York City over the past 400 years we explored the patterns of distant relatedness reflecting fine-scale structure in Bio*Me*. We first detected pairwise genomic tracts inherited IBD (pairwise haplotypic tracts 3cM or longer) between all participants, and 2504 participants comprising 26 global populations of the 1000 Genomes Project Phase 3^25^. We used this information to construct a network of pairwise IBD sharing between all individuals who were not inferred to be directly related (**see methods; Supplementary Figure 3**). To identify ‘communities’ of individuals enriched for recent, shared genealogical ancestry we performed community detection via flow-based clustering using the *InfoMap* algorithm^26,27^ (**Figure 3A** and **3B**).

**Figure 3.**
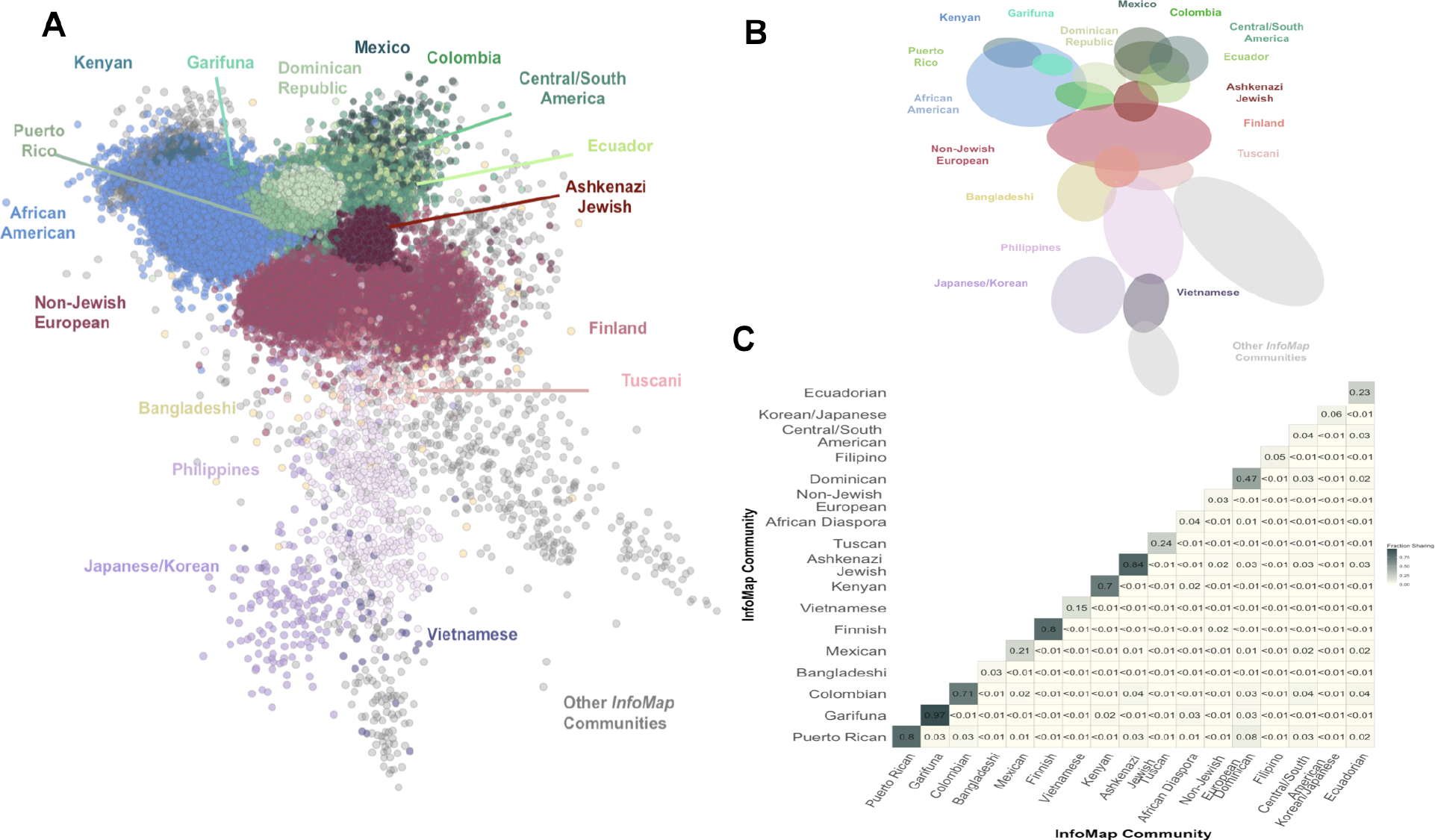
(**A**) Network of IBD sharing among Bio*Me* participants, coloured by community membership as inferred by *InfoMap* (for the top 17 communities only, and only showing nodes with >30 connections to other nodes). Returned communities reflect an enrichment of IBD sharing among individuals of recent, shared ancestry, and thus recapitulate fine-scale population substructure. (**B**) Schematic representation of the 17 distinct IBD communities recovered by *InfoMap*. (**C**) Heatmap representing the population-level fraction of IBD sharing within and between inferred IBD communities reveals a high degree of modularity.

We observed that 96% of Bio*Me* participants fall into one of 17 distinct clusters containing at least 100 individuals that we refer to as IBD communities. Analysis of the inter- and intra-population level sharing of these communities revealed a higher probability of pairwise IBD sharing within compared to between communities (**Figure 3C**). Additionally, in instances where IBD sharing does occur, the pairwise sum of IBD sharing is also higher within than between communities (**Supplementary Figure 4**). We examined the topology of the IBD network and determined community assignments to be non-random through a node-level comparison of similar edges between 10 instances of a network based on pairwise IBD sharing and 100 instances of topologically similar random networks created with the Erdős–Rényi (ER) model^28^. The quantile of the Jaccard similarity coefficient obtained by comparing these networks corresponds to a permutation p-value. Therefore, the communities detected by using the InfoMap algorithm are non-random and statistically significant (empirical permutation p-value< 3.25×10^−4^).

We hypothesized that these communities represent geographical or ethnic substructure within the Bio*Me* population. To test this, we constructed confusion matrices for membership of each community compared to survey-derived population level information for Bio*Me* participants, and reported the Positive Predictive Values (PPV) for each population label versus each identified community (**Supplementary Figure 5**). Many communities had high PPVs (>0.9) with a single country of origin (6/17), including Puerto Rico, the Dominican Republic, Colombia, Ecuador, Mexico and Ethiopia, reflecting recent migrations to NYC. However, a number did not, including one community that notably consisted of 85% of individuals in Bio*Me* who self-reported having Jewish ancestry in the survey (**Supplementary Figure 6**). Notably, other IBD communities were also detected that transcended self-reported ethnicity labels and mapped across various different groups. For example, one community, who, based on a combination of self-reported country-of-origin information and PCA analysis, we suspect are likely to represent Garifuna. Individuals in this community (N=113) were either born in Europe, Central America or in the United States, and self-identified as AA, H/L or Other, but cluster together tightly in principal component space (**Supplementary Figure 7**).

We next determined whether IBD community detection was better able to classify individuals based on recent genetic ancestry than current best practices, *i.e.* k-means clustering over PCA eigenvectors. We performed k-means clustering from k=5 to k=20 over the first 5 principal components calculated across all Bio*Me* participants, and compared k-means clustering to IBD community detection for 6 recent diaspora communities in NYC. To measure accuracy, we calculated PPV, Negative Predictive Value, (NPV), sensitivity and specificity for each inferred cluster using country of origin labels. For the Puerto Rican, Dominican, Ecuadorian, Colombian, Mexican and Ethiopian communities, we observed that PPV, NPV, and specificity was always the same or higher in IBD communities compared to k-means clustering with any value of k (**Supplementary Figure 8**). We noted that, while sensitivity can also be higher for IBD communities, in some cases it was lower than k-means clustering at low *ks* for Colombian, Mexican and Ethiopian groups. We speculate that this may reflect population substructure within a country-of-origin or subsequent patterns of migration to NYC.

### Signatures of Founder Effects in BioMe communities

We observed evidence of founder effects in the form of elevated IBD sharing in multiple IBD communities (**Figure 4A**). These included the Ashkenazi Jewish community (sample size N=4415, median pairwise sum of IBD sharing=21.79cM (95% confidence interval (CI) 21.77-21.80), sum of ROH=10.00MB (95% CI 9.66-10.38), and the Finnish community (N=120, IBD=10.61cM (10.34-10.91), ROH=6.68MB (3.95-7.98)), both of which are known and well-studied founder populations. Five other communities exhibited high levels of IBD sharing (IBD>8cM) and enrichment for autozygosity (ROH>5MB), namely, Puerto Rican (N=5452), Dominican (N=1971), Garifuna (N=113), Colombian (N=234), and Ecuadorian (N=438) communities. In contrast, communities similar to well-studied, non-founder populations like non-Jewish EA exhibited lower levels of IBD sharing (IBD<5cM) and autozygosity (ROH<5MB; **Supplementary Table 3**). Examining the network topology, IBD communities corresponding to founder populations exhibit specific characteristics: they have a high clustering coefficient (e.g. *C* = 0.92 for Ashkenazi Jews; *C* = 0.97 for Garifuna, *C* = 0.85 for Puerto Ricans), while non-founder populations have much lower values (*e.g. C* = 0.05 for AA, *C* = 0.09 for non-Jewish EAs), and they exhibit a strongly bimodal degree distribution, with a high distance between the intra-community degree distribution and the inter-community distribution as summarized by the Wasserstein metric (e.g. W = 0.82 for AJ; W = 0.96 for Garifuna, W = 0.78 for Puerto Ricans), while non-founder populations have much lower values (e.g. *W* = 0.03 for AA, *W* = 0.03 for non-Jewish Europeans) (**Supplementary Table 4**, **Figure 4B**, **Supplementary Figure 9**).

**Figure 4.**
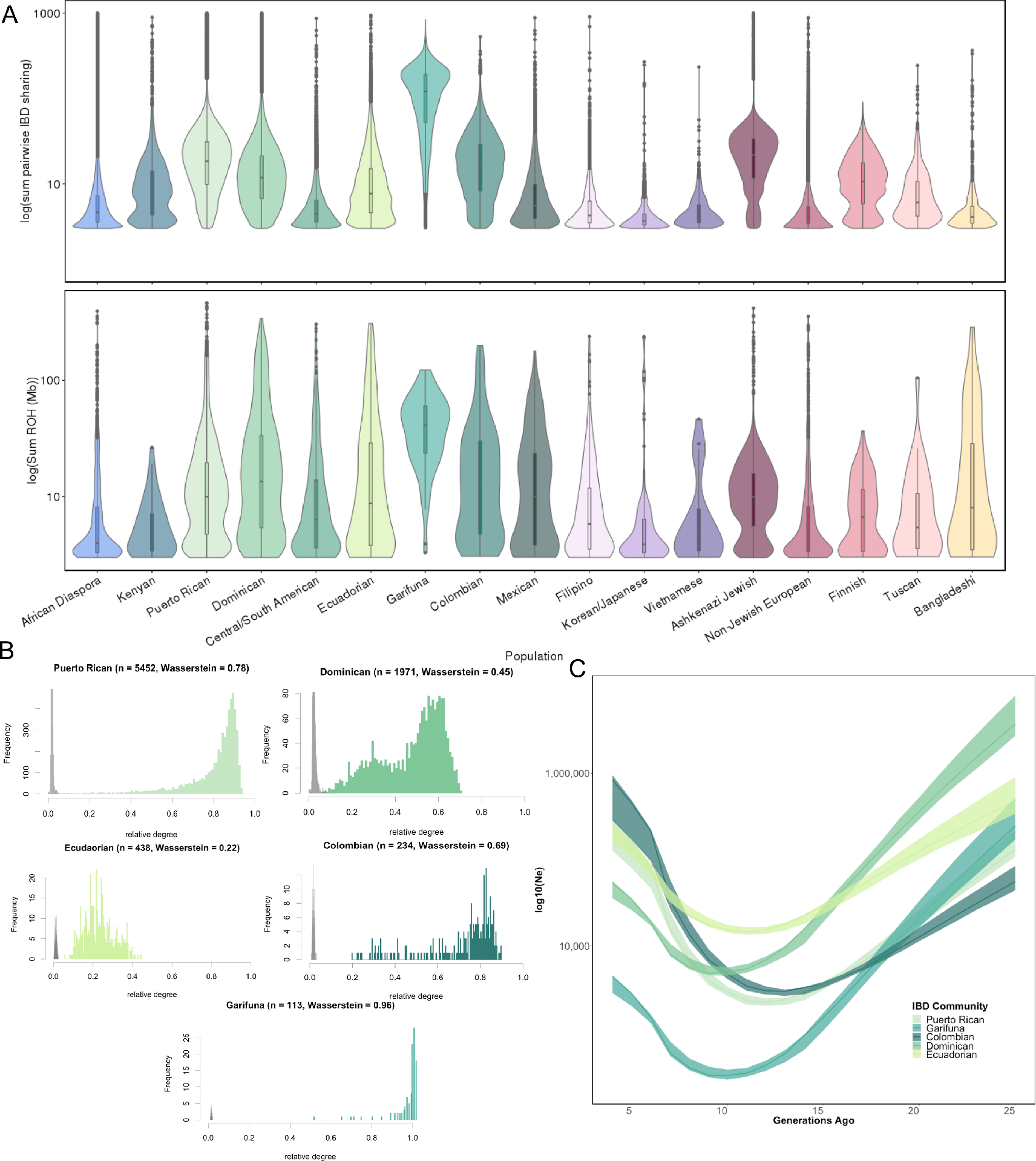
(**A**) (**Top**) Distribution of the mean sum of IBD sharing per inferred IBD-community reveals the presence of elevated IBD sharing in several communities, including canonical founder populations such as the Ashkenazi Jewish and Finnish, but also in several Hispanic/Latino populations including the Puerto Rican, Dominican and Garifuna. (Below) Distribution of the sum of Runs of Homozygosity (ROH) present within individuals per community (**B**) Analysis of the distribution of degree sharing within versus between communities exhibiting elevated IBD revealed high levels of modularity, further suggestive of a founder effect. (**C**) Using the tract length distribution of IBD haplotypes to model the effective population size of communities over antecedent generations revealed evidence of population bottlenecks between 10-15 generations ago.

To understand the timing of the bottlenecks leading to the observed founder effects, we used the IBD_Ne_ software^29^ to model each community’s effective population size (N_e_) over antecedent generations. For the communities representing populations from the Americas, the results are consistent with a population bottleneck occurring approximately 10-15 generations ago, which is coincident with historical accounts of the timings of European contact and subsequent colonization (**Figure 4C**). The most profound bottleneck was observed in the Garifuna community, with a minimum N_e_ of 321 individuals (95% C.I 286-372, 10 generations ago), consistent with previous reports^30^. This analysis demonstrates that founder effects are potentially more ubiquitous than previously thought, with >25% of Bio*Me* participants falling within a community exhibiting elevated IBD sharing, and this was particularly notable among H/L populations.

### Characterizing the Health Phenome in Communities of Distantly Related Individuals

To explore the effect of IBD community membership on predisposition to EHR captured health outcomes, we tested each community for their relative enrichment of health-related traits. First, we extracted ICD-9 (2007 to 2015) and ICD-10 (2015 to present) billing codes from the EHR. We then applied an aggregation schema to convert ICD codes into 1764 distinct diseases and traits called phecodes^31^. We systematically performed logistic regression across all phecodes, where the membership of a given community was used as the primary predictor variable, adjusting for age and sex as covariates, and restricting analyses to communities containing >=1000 Bio*Me* participants (N=7) in total (after excluding the 1000 Genomes reference panel). In all, 1,177 of 4,988 phecode associations tested were either significantly enriched or depleted across all 7 of the top communities after Bonferroni correction. Summary statistics for the relative enrichment and depletion of Bonferroni significant phecode associations per each of the largest 7 communities are reported in **Supplementary Table 5**.

We observed patterns of phecode enrichment across communities of shared continental ancestry. For example, 3 associations were significantly enriched in all three communities with appreciable African genetic ancestry (>20%), namely the Puerto Rican, Dominican, and African diaspora communities (**Supplementary Table 5**). These codes include essential hypertension, peripheral vascular disease and type 2 diabetes (**Supplementary Figure 11**). Higher prevalence of these diseases within the African American populations have been widely reported in the epidemiological literature, but our findings, and emerging evidence, support these diseases as being prevalent in some Hispanic/Latino communities too.

Of the significantly associated phecodes, 21.7% (n=274) were uniquely enriched in a single community, many of which had been previously reported as being at increased incidence within those communities. For example, the phecode for asthma was observed to be most highly enriched in the Puerto Rican ancestry community (**Supplementary Figure 11**; phecode=495; O.R.=2.91 (95% C.I.=2.70-3.14); p<1.126×10^−169^). Further, we observe elevated levels of ulcerative colitis (phecode=555.2; O.R=2.61 (1.99-3.42); p<2.904×10^−12^) and Parkinson’s disease (phecode=332; O.R=2.31 (1.78-3.00); p<4.308×10^−10^) in the Ashkenazi Jewish community (**Supplementary Figure 11)**. Within the non-Jewish European ancestry (**Supplementary Figure 11**) community we observe elevated levels of “multiple sclerosis” (phecode=”335”; 2.55 (2.01-3.237); p <1.328×10^−14^) and “Basal cell carcinoma” (phecode=”172.21; 3.24 (2.50-4.20); p <7.845×10^−19^). For the Filipino community, both viral hepatitis B (phecode=”560”, 6.60 (5.01-8.69); p< 3.9×10^−41^) and gout (phecode=“363” 2.94 (1.99-4.35); p < 6.551×10^−08^) both appeared to be significantly enriched (**Supplementary Figure 11**).

Among the community specific phecodes, we also observe an enrichment of phecodes suggesting increased prevalence of diseases within communities that had not been previously reported. We observed a cluster of significantly enriched circulatory system (CS)-related phecodes, where 82% (9/11) were significantly enriched in the Dominican community. For example atherosclerosis of native arteries of the extremities (phecode=“440.22”; 4.06 (3.13-5.28); p<9.83×10^−26^) and coronary atherosclerosis (phecode=“411.4”; 1.64 (1.43-1.88); p<1.61×10^−12^) were the top two most significantly enriched codes (**Figure 5E**), suggestive of underlying increased prevalence of peripheral artery disease (PAD) in this population. Likewise, we observed a cluster of significantly enriched endocrine and metabolic-related phecodes, with 28% (9/32) significantly enriched in Ashkenazi Jews. The top phecodes are chronic lymphocytic thyroiditis (phecode=“245.21”; 4.16 (3.62-4.78); p<2.31×10^−89^) and hypothyroidism (phecode=“244.4”; 1.65 (1.49-1.83); p<3.0×10^−21^), suggesting an increased prevalence of autoimmune thyroid disease in this population. Taken together, this analysis suggests that identification of fine-scale genetic communities can help to elucidate the presence of population health disparities within healthcare systems.

**Figure 5.**
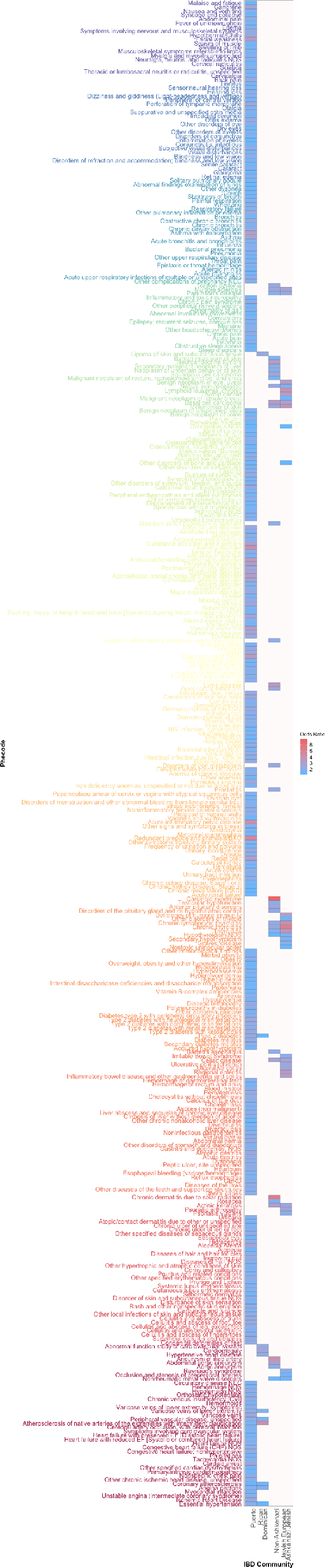
Heatmap of significantly enriched Phecodes in the Puerto Rican, Dominican, Ashkenazi Jewish, and Non-Jewish European IBD communities reveals patterns of fine-scale population health disparities.

## Discussion

Improving health outcomes in populations around the world will require strategies for earlier detection and better prevention of disease. Achieving this goal will necessitate greater precision in our health systems so that the right intervention can be applied in the right population at the right time. Recent advances in genomic technology have resulted in a rapid increase in the amount of human genomic data in health systems^6^. Here we demonstrate how this data may be used for fine-scale population health monitoring. We show how poorly demography information is captured in a large urban health system, particularly for non-European populations. We apply a framework to detect fine-scale population structure by characterizing a network of distant relatedness within patients in the Bio*Me* biobank, and detect 17 distinct communities that are highly correlated with culturally endogamous groups and recent diaspora to New York City from countries around the world. We demonstrate that IBD community detection robustly and stably recapitulates recent patterns of demography in NYC, and similar ideas have been explored in previous work^32–34^. By linking to Electronic Health Records and testing for enrichment of health outcomes within uncovered communities, using phenotypes derived from ICD-9 and ICD-10 billing codes, we demonstrate a significant community-specific enrichment of both anticipated and novel health related traits.

We show that many of the community’s exhibit evidence of founder effects, as demonstrated by elevated levels of autozygosity, median within-community IBD sharing, and by applying methods that measure the degree of distance between communities in network topology. This includes canonical founder populations with known historical evidence of founder events, including Ashkenazi Jewish^35^, Finnish^33^ and Garifuna populations^35,36^. We also show evidence for the timing and magnitude of founder effects in less characterized H/L founder populations in NYC, namely populations of Puerto Rican, Colombian, Ecuadorian, and Dominican descent. We demonstrate these founder effect occurred 10-14 generations ago, coincident with the era of colonization of the Americas. This finding is supported by previous work from our group and others demonstrating evidence of founder effects in populations from the Caribbean and Americas^24,37,30^, and reviewed here^38^. A better understanding of founder effects in Hispanic/Latino populations may help uncover monogenic disease variants segregating at appreciable frequency, for example the recently discovered variant underlying Steel Syndrome in Puerto Rican populations. This indicates an under recognized potential for reducing the genetically driven health burden within Hispanic/Latino communities, echoing similar work in South Asian populations^39^.

Overall approximately one quarter of Bio*Me* participants harbor genetic signatures of founder effects, and extrapolating this observation to NYC, we estimate that approximately 15% of New Yorkers can be genetically linked to one or more founder populations. This finding mirrors similar observations of founder effects in large, predominantly European ancestry biobanks in Finland^33^ and rural Pennsylvania^40^, where founder effects we found to be ubiquitous in the former and in 22% of the latter cohort. A recent study of a direct-to-consumer genetic database of approximately 770,000 customers across the US also revealed myriad signatures of founder effects which could be attributed to both pre-diaspora population structure and/or post-diaspora isolation, i.e. multiple Irish ancestry groups in Boston^32^. This suggests that founder effects and founder populations may be more ubiquitous that previously thought, and that as yet little understood processes of diaspora and migration can contribute to these effects. These findings are clinically important, as communities of recently related individuals may be more likely to share rare, clinically relevant variants. For example, we recently showed that 50% of carriers of pathogenic variants in *BRCA1* and *BRCA2* genes that are linked to hereditary breast and ovarian cancer syndrome in Bio*Me* are carrying known founder mutations. Finally, the network-based machine learning approaches applied here are highly scalable to very large datasets of individuals ascertained through unselected, population-based recruitment processes. As more and more genomic data becomes available, via projects like the UKBiobank and the All of Us Research Program, we anticipate approaches like ours will uncover increasing nuances in the population structure inherent in these databases.

The fine-scale population structure revealed by the IBD communities can also have important implications for genomic medicine. By linking an individual’s community membership with EHRs, we were able to use a phenome-wide approach to investigate enrichment and depletion of diseases in 7 of the largest IBD communities. In many cases these associations recapitulate known disease prevalence in specific populations, for example increased rates of gout and viral hepatitis B in the Filipino community. However, the comparison of associations across and within communities can also expand our current understanding of these prevalence, for example, showing that chronic kidney disease not only impacts African-Americans, but also Puerto Rican and Dominican populations in NYC. Finally, this type of fine-scale population analysis can also reveal previously under-appreciated health risks impacting specific populations. We demonstrated evidence for enrichment of suspected peripheral artery disease in Dominican and autoimmune thyroid disease in Ashkenazi Jews, and future work should investigate factors, including genetic, lifestyle, environmental and health care access, which might impact these outcomes. Increasing resolution of longitudinal phenome data in health systems will offer enhanced opportunities to monitor health in real-time, allowing for more agile programs for discovery, prediction and intervention.

## Materials and Methods

### Ascertainment of Ethnicity Information in BioMe

Information regarding participant’s ethnicity was ascertained as part of the BioMe questionnaire which is completed at enrollment. Participant ethnicity was solicited in the form of the multiple-choice question with 9 options to choose from (**Supplementary Table 6**). Prior to 2014, in addition to a multiple-choice question about ethnicity, participants were also given the option to report their country of birth. After 2014, enrolling participants were provided with options to report the country of birth of both their parents and grandparents as well. All participants recorded answers to survey question 1, and 43.6% of participants also provided responses to survey question 2.

### Comparison of Electronic Health Record versus self-reported Ethnicity

Ethnicity information was extracted from the Electronic Health Records (EHR) for all BioMe participants for every available patient visit between January 2007 and December 2014. The possible ethnicity designations within this timeframe consisted of “African American (Black)”, “Asian”, “Caucasian (White)”, “Hispanic/Latino”, “Native American”, “Other”, “Pacific Islander” or “Unknown”.

For individuals who had greater than one interaction with the healthcare system and conflicting EHR recorded ethnicity, we selected the ethnicity designation assigned at their earliest visit for downstream comparison to self-reported ethnicity. We mapped EHR recorded ethnicity to self-reported ethnicity for N=36061 Bio*Me* participants across 1492428 healthcare visits in total.

For individuals who only self-reported one ethnicity, in instances where that category had a direct mapping to one of the EHR ethnicity variables, we made a direct comparison between self-reported and EHR recorded ethnicity to calculate the percentage of individuals who had been miss-classified in the EHR. With the exception that we mapped individuals who self-identified as “Caucasian/White” (N=7691), “Jewish” (N=934), or both (N=935) to the EHR category of “Caucasian (White)”. We also mapped individuals who self-identified as *“East or Southeast Asian (i.e. China Japan Korea Indonesia)”* only to the EHR category “Asian”, while individuals who self-identified as *“South Asian/Indian (i.e. India Pakistan)“* were analyzed separately.

### Genotype Quality Control

BioMe participants (N=32595) were genotyped on the Illumina Global Screening Array (GSA) platform. Quality control of the GSA data for N=32595 participants and n=635623 variants was performed stratified by ethnicity category. Individuals with an ethnicity-specific heterozygosity rate that surpassed +/− 6 standard deviations of the population-specific mean, along with individuals with a call rate of <95% were removed (N=684 participants in total). N=80 individuals were then removed for exhibiting persistent discordance between EHR-recorded and genetic sex. A further N=126 duplicate individuals were also excluded from downstream analysis. In total 31,705 passed sample level QC for downstream analysis. All quality control steps were conducted using Plink(v1.90b3.43)^41,42^. Sites with a call rate below 95% were excluded (n=19253), along with sites that were seen to significantly violate Hardy-Weinberg equilibrium (HWE) when calculated stratified by ancestry. HWE thresholds for site exclusion varied by ethnicity, specifically we set a threshold of p < 1e-5 in for all populations except Hispanic/Latinos’ where it was set to p < 1e-13 (n=11503 SNPs in total). This resulted in the retention of n=604869 sites.

### Genetic Relatedness Estimation

Pairwise kinship coefficients were estimated for all Bio*Me* participants (N=31705) using all N=604869 SNPs that passed QC using the KING software^43^ (v1.4) by passing the --*kinship* flag.

### Global Ancestry Estimation

Prior to calculating PCA we restricted analysis to common (minor allele frequency (MAF) >0.01), autosomal sites. We also removed regions of the genome known to be under recent selection, specifically *HLA* (chr6: 27032221-35032223, hg38), *LCT* (chr2:134242429-136242430), an inversion on chromosome 8 (chr8:6142478-16142491), a region of extended LD on chromosome 17 (chr17:41843748-46922634), *EDAR* (chr2:108383544-109383544), *SLC2A5* (chr15:47707803-48707803), and *TRBV9* (chr7:142391891-142392412). Finally, the GSA data was intersected with and merged with genome sequence data from 26 populations in 1000 Genomes Project phase 3 reference panels (TGP; N=2504), and 53 populations in the Human Genome Diversity Panel^44^ (HGDP; N=986) and 8 additional reference panels (Bari, Khomani, Nama, Oaxacan, Peru Warao, Yukpa, Zapotec) from the Population Architecture using Genomics and Epidemiology (PAGE; N=700) study, both genotyped on the Multi-Ethnic Genotyping Array (MEGA). This resulted in a total of n=260502 snps and N=35854 individuals. The first 20 PCs were calculated using PLINK (v1.9). We also ran ADMIXTURE^19^ with 5-fold cross validation from *k*=2 to *k*=12 across all individuals inferred to be unrelated (N=32354), by randomly removing one of each individual in a pairwise relationship defined by KING to be greater than 3^rd^ degree relatives (as defined by a pairwise kinship coefficient of >= 0.0442 in the KING output). To visualize fine-scale population substructure we applied the Uniform Manifold Approximation and Projection (UMAP) to the first 10 principal components across all samples using the “umap” library in R using the default parameters.

### Phasing and Identity-by-Descent Inference

Prior to phasing and inference of Identity-by-Descent, the GSA array data for Bio*Me* participants was lifted over to GRCh37/hg19, before additional quality control was performed on variants, including the removal of SNPs with a call rate below 99% (n=48436) and variants with a MAF <1% (n=135011). Palindromic variants were also excluded at this stage (n=4375). The data was subsequently merged with the TGP reference panels (N=2504 individuals), and only intersecting sites were retained, resulting in the retention of n=402042 SNPs in total. Phasing was subsequently performed per autosome on all N=34209 individuals with the Shapeit software^45^(*v2.r790*) using the hapmapII genetic map (build: GRCh37/hg19) using default flags and – output-max –force. Phased haplotypes were subsequently converted to plink format and IBD was called using the iLASH software using the following flags:

*slice_size 400*, *step_size 400*, *perm_count 12*,*shingle_size 20*, *shingle_overlap 0*, *bucket_count 4*, *max_thread 20*, *match_threshold 0.99*, *interest_threshold 0.70*, *max_error 0*, *min_length 3*

For quality control, IBD tracts that overlapped with low complexity regions were excluded, along with IBD tracts that fell within regions of excessive IBD sharing, which we defined as regions of the genome where the level of pairwise IBD sharing exceeded 3 standard deviations above the genome-wide mean **(Supplementary Figure 10**).

### Network Construction and Community Detection

To construct the IBD network, IBD tracts along the genome were summed between each pair of individuals inferred to be 2^nd^ degree relatives or less from Bio*Me* and the 1000 genomes reference panel (N=31683 individuals in total) to generate the total sum of IBD sharing per pairwise relationship. This was used to construct an adjacency matrix where each node represents an individual and each weighted edge represents the pairwise sum of IBD sharing. To detect the presence of structure within the IBD network we used the implementation of InfoMap in the iGraph package (R version 3.2.0). Visualization of the IBD network was also performed using iGraph using a Fruchtermen-Reingold layout (with n=1000 iterations), after excluding poorly connected nodes (< 30 connections).

Community assignments were ascertained by using InfoMap both on the original IBD-based network (10 times) and on 100 sets of topologically similar random networks generated with the ER model. We defined a neighborhood matrix as a n × n matrix where Mi,j = 1 if samples i and j are part of the same community and = 0 if they aren’t. For an individual k, the Jaccard similarity coefficient obtained while comparing the k-th row of two neighborhood matrices constitutes a natural measure of community relatedness, with J = 1 if k has exactly the same neighbors in the two matrices, J = 0 if k does not share any neighbors between the two matrices, and 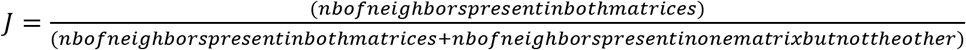. Thereby, an empirical p-value can be obtained for each individual, by comparing the Jaccard coefficient obtained with neighborhood matrices derived from the original IBD-based network and the distribution of Jaccard coefficients obtained with a matrix derived from the original network and a matrix derived from a random network. Such an empirical p-value can be extended to the whole network comprising all community assignments by computing the Jaccard coefficient with the complete neighborhood matrices. To ascertain scale-freeness, we computed the degree distributions of each community: for each node of the network, we count the number of edges connecting to another node assigned to the same community by infomap (intra-community degree) and the number of edges connecting to another node assigned another community (inter-community degree). For comparison purposes, degrees are normalized by n−1, where n is either the community size in the case of intra-community degree or the sum of all the other communities’ sizes in the case of the inter-community degree; such that an individual sharing IBD with all the other individuals from the same community has a relative within-community degree of 1, and an individual sharing IBD with all the other individuals from all the other communities had an out-of-community degree of 1. To ascertain small-world properties, we computed the average shortest path length between two nodes and the average clustering coefficient (the ratio of existing edges between the neighbors of a specific node relative to the total number of potential edges between said neighbors) and compared them with the same values obtained from an ER random network^46^. To quantify bimodality and distance between inter- and intra-community degree distributions, we used the Wasserstein distance^47^.

### Analysis of IBD Community Membership Using Population Labels

To explore the correlation between self-reported country of origin, subcontinent and ethnicity and IBD community membership for Bio*Me* participants we calculated Positive Predictive Values (PPVs), Negative Predictive Values (NPVs), Sensitivity and Specificity for each self-reported label versus membership of each IBD community using the “caret” library in R^48^. For the analysis by country and subcontinental origin, US-born Bio*Me* participants were excluded from the calculations.

### Inference of Runs of Homozygosity

Genotype data for all individuals included in the network analysis (N=31683) was LD thinned using PLINK(v1.90b3.43) using the “*--indep-pairwise 50 5 10*” command. This resulted in the retention of n=477,342 SNPs in total. Runs of homozygosity were then calculated using the following commands: “*--homozyg-density 200*, *--homozyg-gap 500*, *--homozyg-kb 3000*, *--homozyg-snp 65*, *--homozyg-window-het 0*, *--homozyg-window-missing 3*, *--homozyg-window-snp 65*”.

### Phenotype ontology

15,665 unique ICD-9 and ICD-10 billing codes from the Mount Sinai Bio*Me* biobank were collapsed into 1764 phenotype codes (Phecodes)^31,49^ on the basis of the PheWAS catalog (*www.phewascatalog.org*) using an in-house R script (R version 3.4.1).

### Analysis of Enrichment of Phecodes within IBD-Communities

We performed logistic regression systematically across all 1764 phecodes and all IBD communities with >=1000 members who were also Bio*Me* participants (N=7 in total), (*i.e.* excluding communities predominantly composed of individuals from the 1000 Genomes Reference panel). To do so we encoded IBD community membership as a binary predictor variable and generated Plink format “ped” files for each community, where community membership was encoded as “1” and non-membership encoded as “2”. We then performed logistic regression for each phecode and for each community using using Plink(v1.90b3.43), adjusting for age and sex as covariates, and excluding 2nd degree relatives and above. To avoid spurious association, phecodes that were present in less than 10 instances per community were excluded from the analyses. Statistical significance was determined for each IBD community-wide association via Bonferroni correction.

