## Supplemental Information for "Towards a fine-scale population health monitoring system"

**CBIPM Leadership Team**

Noura Abul-Husn, Erwin Bottinger, Judy Cho, Ron Do, Steve Ellis, Omri Gottesman, Yuval Itan, Eimear Kenny, Ruth Loos, Amanda Merkelson, Girish Nadkarni, Aniwaa Owusu-Obeng.

Contribution: All authors contributed to securing funding, study design and oversight.

**Sequencing and Lab Operations**

Bernadette Liggayu, Amanda Merkelson, Janice Morinigo, Patrick Shanley, Quingbin Song

Contribution: All authors are responsible for DNA extraction, sample handling and tracking, and the library information management system.

**Clinical Informatics**

Noura Abul-Husn, Lili Chan, Steve Ellis, Omri Gottesman, Arden Moscati, Girish Nadkarni, Rajiv Nadukuru, Aniwaa Owusu-Obeng, Teilman Van Vlick

Contribution: All authors are responsible for analysis needed to produce electronic health record extracted data.

**Genome Informatics**

Gillian Belbin, Dean Bobo, Kumardeep Chaudhary, Sinead Cullina, Ron Do, Aine Duffy, Amanda Dobbyn, Yuval Itan, Eimear Kenny, Margret Linan, Carla Marquez-luna, Arden Moscati, Ha My, Micheal Preuss, Cigdem Sevim, Stephane Wenric, Ryan Walker, Zhe Wang, Yiming Wu.

Contribution: All authors are responsible for analysis needed to produce exome and genotype data.

**Regeneron Genetics Center Banner Author List and Contribution Statements**

All authors/contributors are listed in alphabetical order.

**RGC Management and Leadership Team**

Goncalo Abecasis, Aris Baras, Michael Cantor, Giovanni Coppola, Aris Economides, John D. Overton, Jeffrey G. Reid, Alan Shuldiner.

Contribution: All authors contributed to securing funding, study design and oversight. All authors reviewed the final version of the manuscript.

**Sequencing and Lab Operations**

Christina Beechert, Caitlin Forsythe, Erin D. Fuller, Zhenhua Gu, Michael Lattari, Alexander Lopez, John D. Overton, Thomas D. Schleicher, Maria Sotiropoulos Padilla, Karina Toledo, Louis Widom, Sarah E. Wolf, Manasi Pradhan, Kia Manoochehri, Ricardo H. Ulloa.

Contribution: C.B., C.F., K.T., A.L., and J.D.O. performed and are responsible for sample genotyping. C.B, C.F., E.D.F., M.L., M.S.P., K.T., L.W., S.E.W., A.L., and J.D.O. performed and are responsible for exome sequencing. T.D.S., Z.G., A.L., and J.D.O. conceived and are responsible for laboratory automation. M.P., K.M., R.U., and J.D.O are responsible for sample tracking and the library information management system.

**Genome Informatics**

Xiaodong Bai, Suganthi Balasubramanian, Leland Barnard, Andrew Blumenfeld, Yating Chai, Gisu Eom, Lukas Habegger, Young Hahn, Alicia Hawes, Shareef Khalid, Jeffrey G. Reid, Evan K. Maxwell, John Penn, Jeffrey C. Staples, Ashish Yadav.

Contribution: X.B., A.H., Y.C., J.P., and J.G.R. performed and are responsible for analysis needed to produce exome and genotype data. G.E., Y.H., and J.G.R. provided compute infrastructure development and operational support. S.K., S.B., and J.G.R. provide variant and gene annotations and their functional interpretation of variants. E.M., L.B., J.S., A.B., A.Y., L.H., J.G.R. conceived and are responsible for creating, developing, and deploying analysis platforms and computational methods for analyzing genomic data.

**Planning, Strategy, and Operations**

Paloma M. Guzzardo, Marcus B. Jones, Lyndon J. Mitnau.

Contribution: All authors contributed to the management and coordination of all research activities, planning and execution. All authors contributed to the review process for the final version of the manuscript.

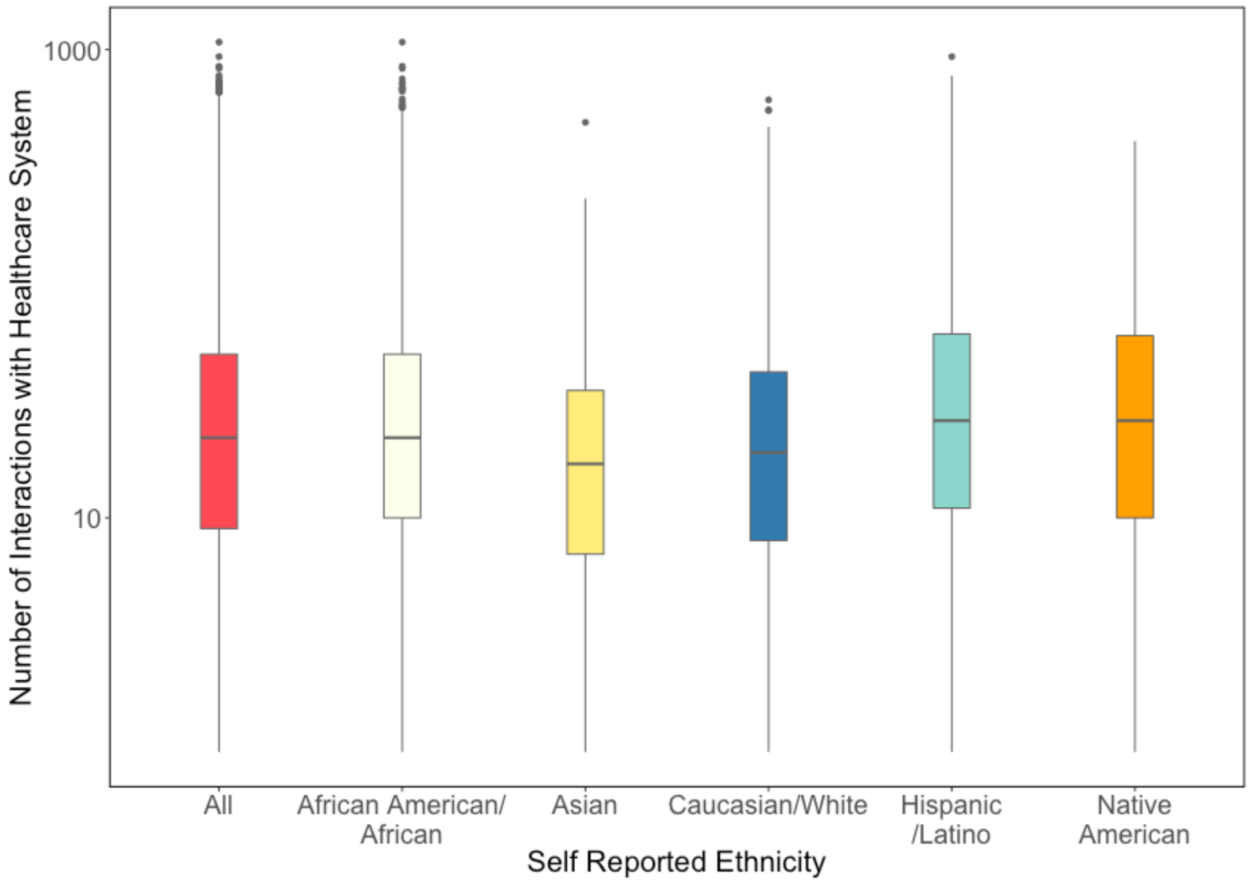

**Supplementary Figure 1. Related to Figure 1.** Distribution of the Number of Interactions with the Mount Sinai Healthcare System Per Individual between 2007-2014, both for Bio*Me* overall, and stratified by self-reported ethnicity.

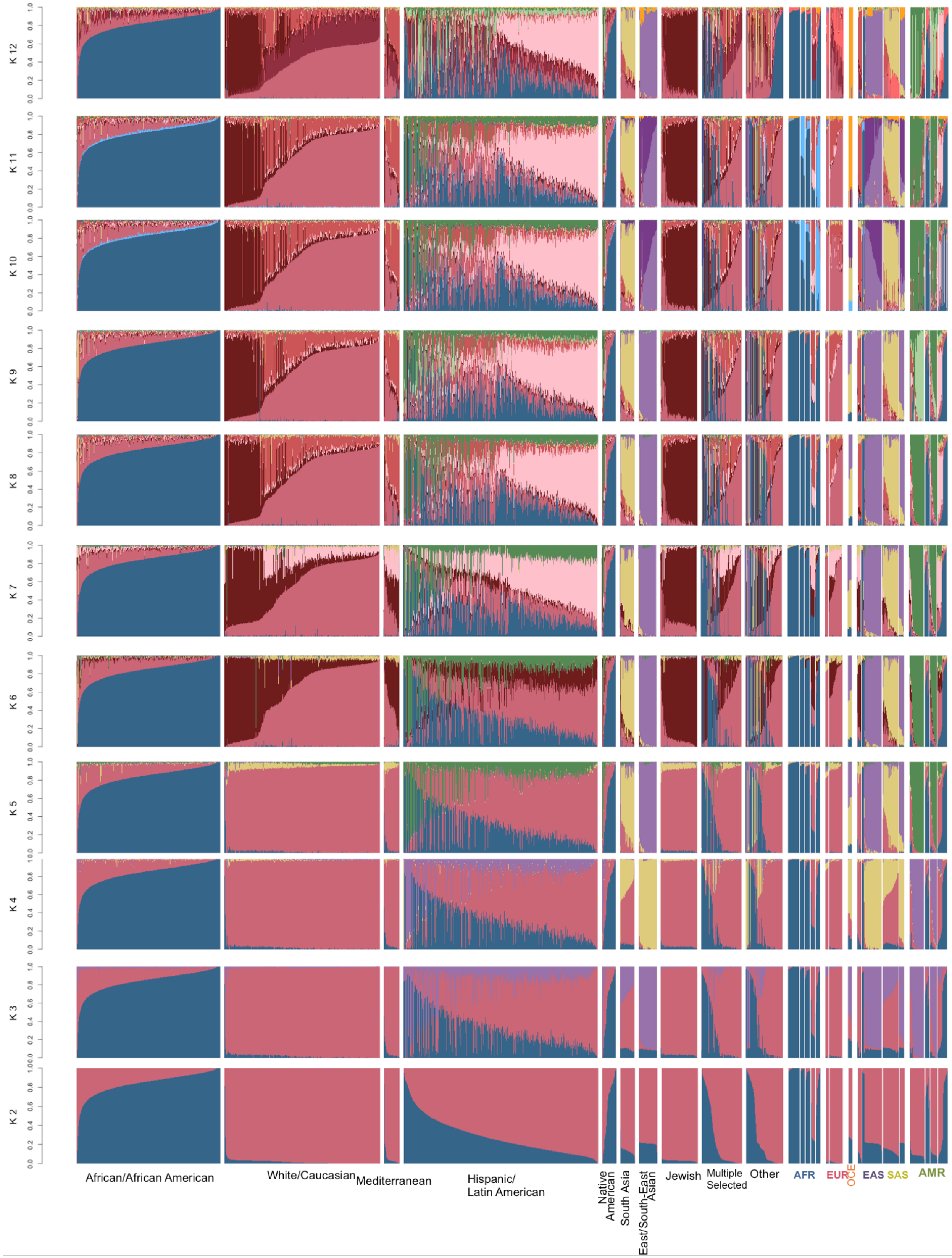

**Supplementary Figure 2. Related to Figure 2. ADMIXTURE runs from k=2 to k=12 in**

**N=31705 unrelated Bio*Me* participants and reference samples from 87 global populations.** ADMIXTURE output of Bio*Me* participants stratified by their self-reported ethnicity (left) and reference samples from 87 global populations (right). Populations labelled “AFR” correspond to reference samples from Africa, “EUR” from Europe, “OCE” from Oceania, “EAS” from East Asia, “SAS” from South Asia and “AMR” from the Americas.

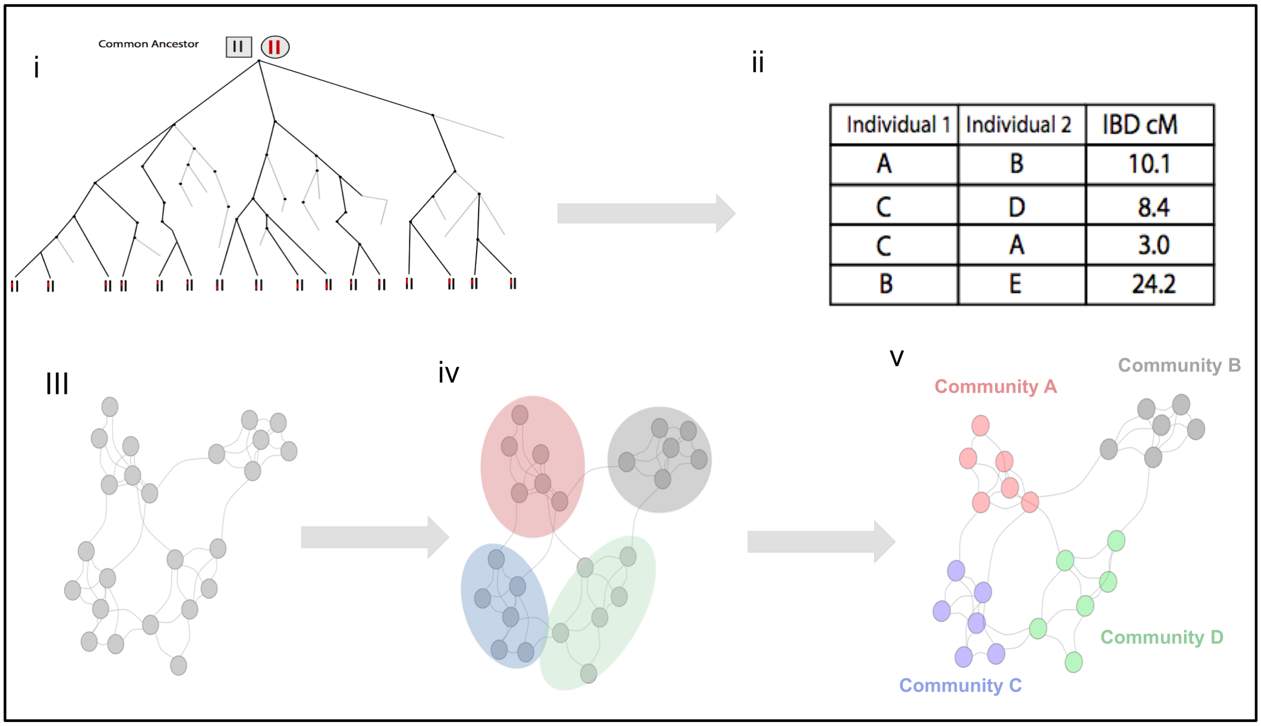

**Supplementary Figure 3**. **Related to Figure 3**. Schematic of the IBD-community detection workflow. (**i**) Haplotypes inherited Identical-by-Descent (IBD) from a recent, common ancestor are present and readily detectable in genomic data. (**ii**) Detected IBD segments can be used to construct an adjacency matrix where the pairwise relationship between each Bio*Me* participant is represented by the total sum of IBD segments they share across their genome (cM). (**iii**) This adjacency matrix can be used to construct a network where every node represents an individual and each (weighted) edge represents the sum of IBD sharing between a given pair. (**iv**) Running the community detection algorithm *infomap* over the IBD network allows for the detection of 'communities' of individuals that are statistically enriched for the sharing of IBD. (**v**) *Infomap* returns community membership status for each node in the network.

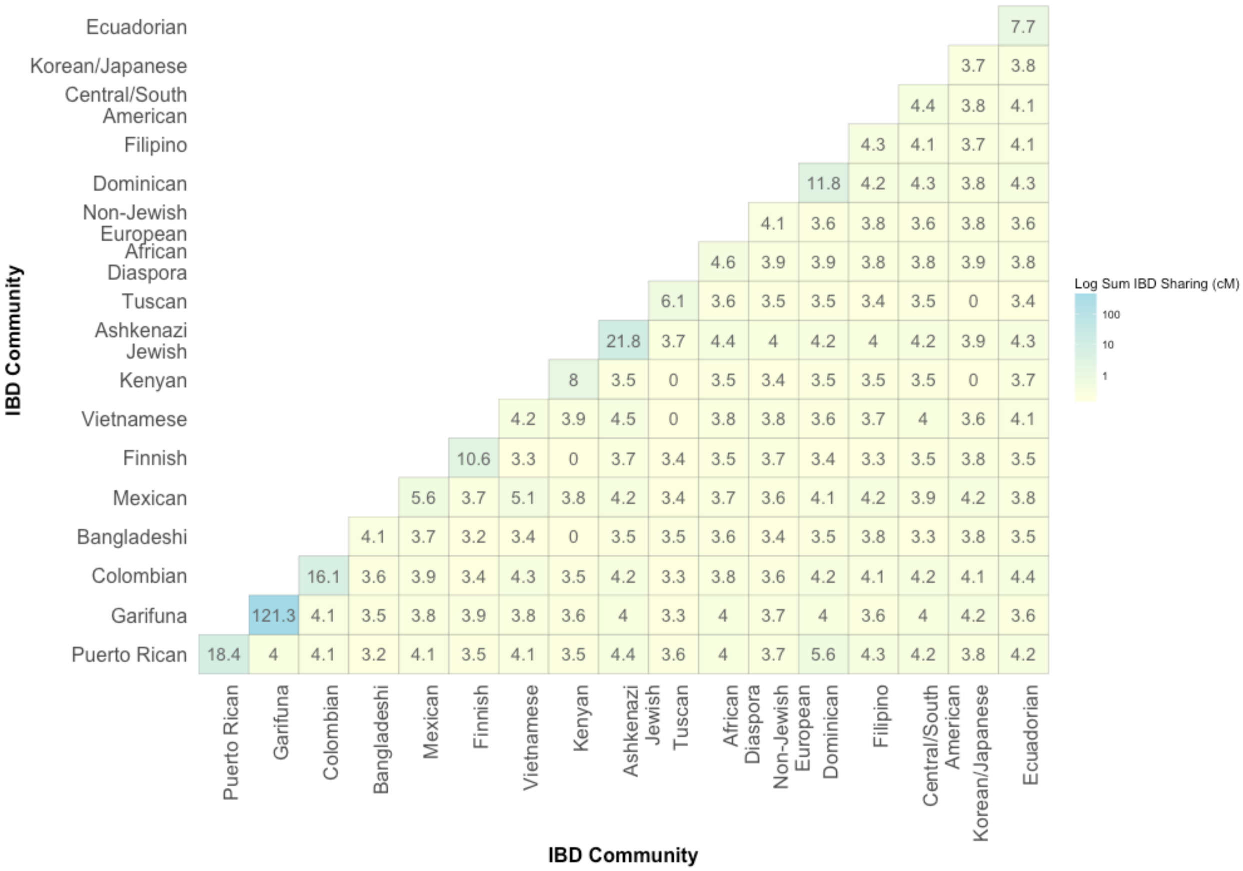

**Supplementary Figure 4. Median sum of IBD sharing within and between the largest IBD Communities in Bio*Me*. Related to Figure 3.** Each tile within the heatmap represents the median sum of IBD haplotype sharing ($\log_{10}$ scale) within and between- IBD communities, with blue representing higher IBD sharing and yellow representing lower levels of sharing.

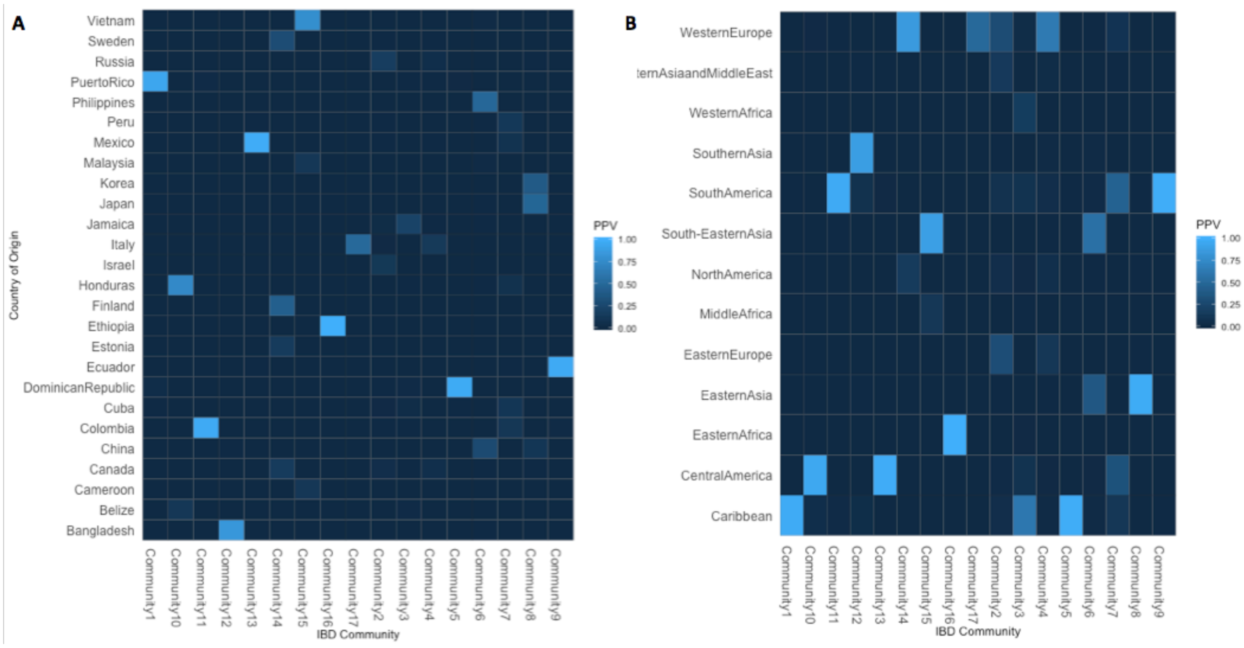

**Supplementary Figure 5. Positive Predictive Values (PPVs) for geographical origin versus IBD Communities. Related to Figure 3. (A)** PPVs for country of origin versus the 17 largest IBD communities. (**B**) PPVs for sub-continental region of origin. Results are only shown for population labels with a PPV >= 0.1 for at least one IBD community.

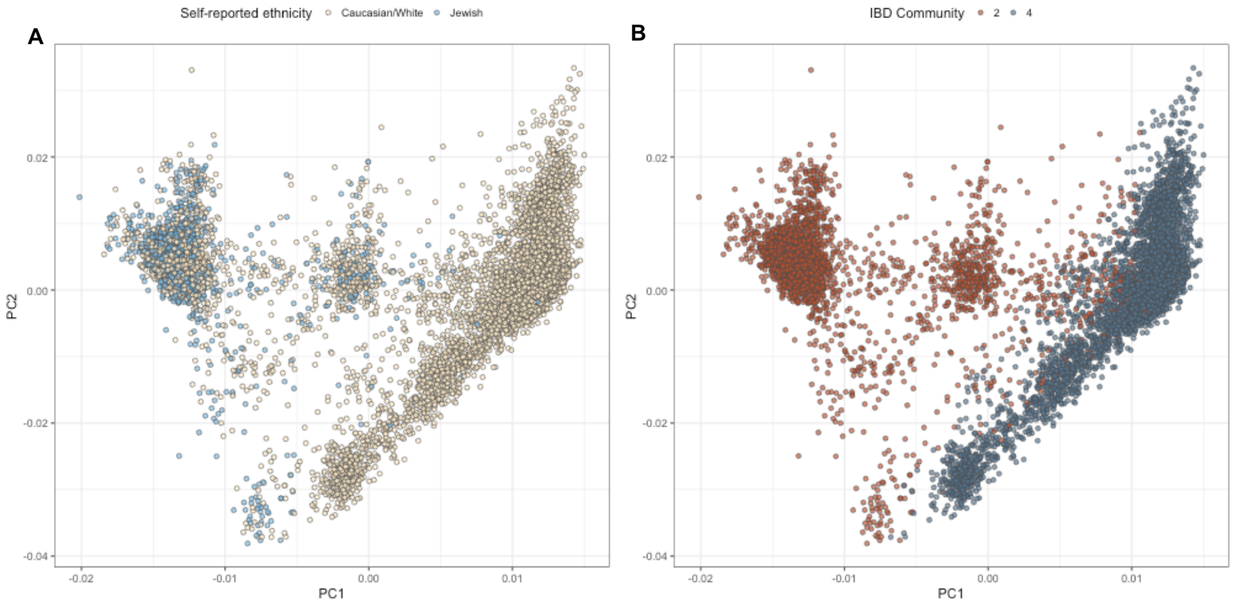

**Supplementary Figure 6. PCA analysis of Bio*Me* European Americans reveals IBD-Community membership reflects genetic Jewish versus non-Jewish ancestry. Related to Figure 3.** (**A**) PCA plot of Bio*Me* participants who self-identify as "Jewish" (blue) and "Caucasian/White" (yellow) reveal clustering in PCA space. (**B**) The same PCA plot coloured by IBD-community membership, where community "2" (red) appears to represent genetic Jewish ancestry, while community "4" (dark blue) represents Non-Jewish European ancestry.

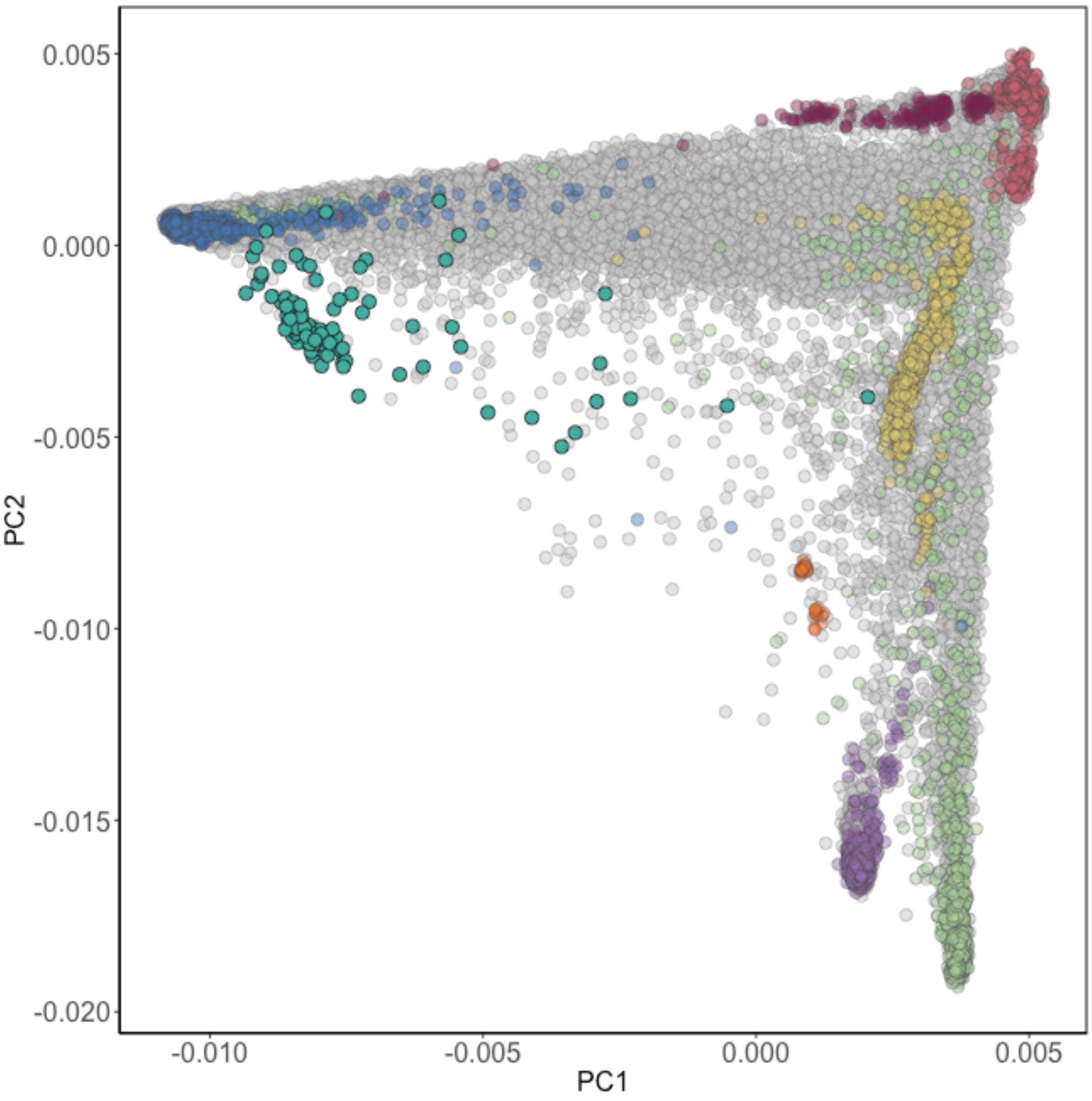

**Supplementary Figure 7. PCA analysis with of the Bio*Me* IBD community inferred to be Garifuna. Related to Figure 3.** Bio*Me* participants who belong to the IBD community that we infer are likely to be Garifuna (teal) cluster on a cline between the African (blue) and Native American (green) reference panels.

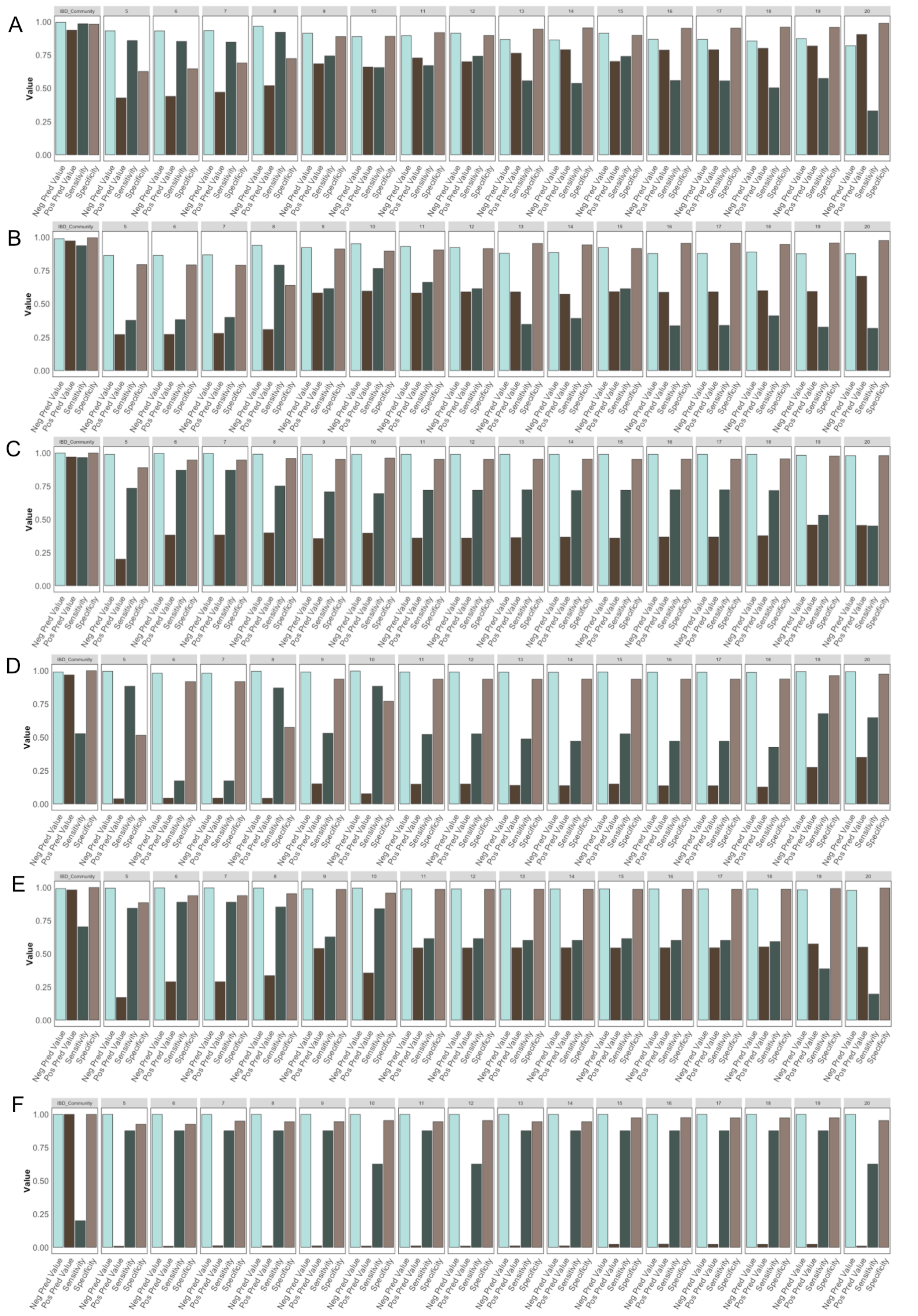
A
B
C
D
E
F
IBD_Community
Value
1.00
0.75
0.50
0.25
0.00
Neg Pred Value
Pos Pred Value
Sensitivity
Specificity

**Supplementary Figure 8. IBD based community detection versus k-means clustering over principal component analysis. Related to Figure 3.** Comparison of population designation using k-means clustering versus IBD communities with strong PPVs (>0.9) for a particular country of origin. The first panel of each plot represents classification metrics for IBD community detection against a given country of origin, specifically Puerto Rican (**A**), Dominican (**B**), Ecuadorian (**C**), Colombian (**D**), Mexican (**E**) and Ethiopian (**F**). The following panels represent the same metrics for the cluster obtained via k-means clustering over PCA with the highest PPV for a given specification of *k* (ranging from k=5 to k=20).In each instance the IBD-community detection is constantly able to better classify individuals in recent diaspora populations based on Positive Predictive Value, Negative Predictive Value, Sensitivity and Specificity.

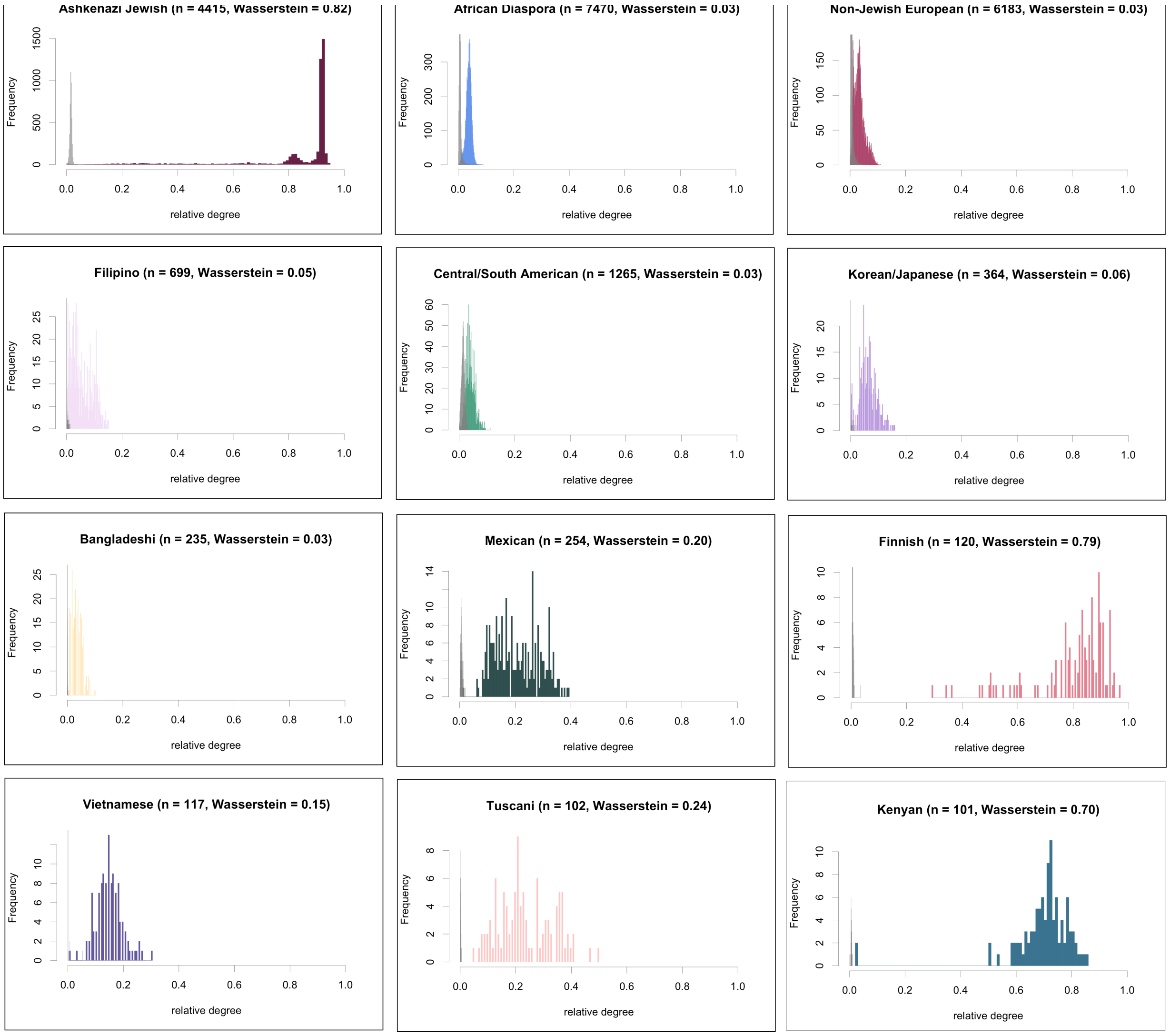

**Supplementary Figure 9. Distribution of the Intra- and Inter- degree distributions within and between IBD communities. Related to Figure 3.** Analysis of the distribution of degree sharing within versus between community for the Ashkenazi Jewish, African Diaspora, Non-Jewish European, Filipino, Central/South American, Korean/Japanese, Bangladeshi, Mexican, Finnish, Vietnamese, Tuscan and Kenyan communities. The within community degree distribution is shown in color and the between communities degree distribution is shown in gray. The strong bimodality in the degree distributions of the Ashkenazi Jewish and Finnish communities, quantified by a Wasserstein metric value of 0.82 and 0.79 respectively, are indicative of a founder effect. The distributions related to the other communities show low bimodality, quantified by Wasserstein metric values ranging from 0.03 to 0.24

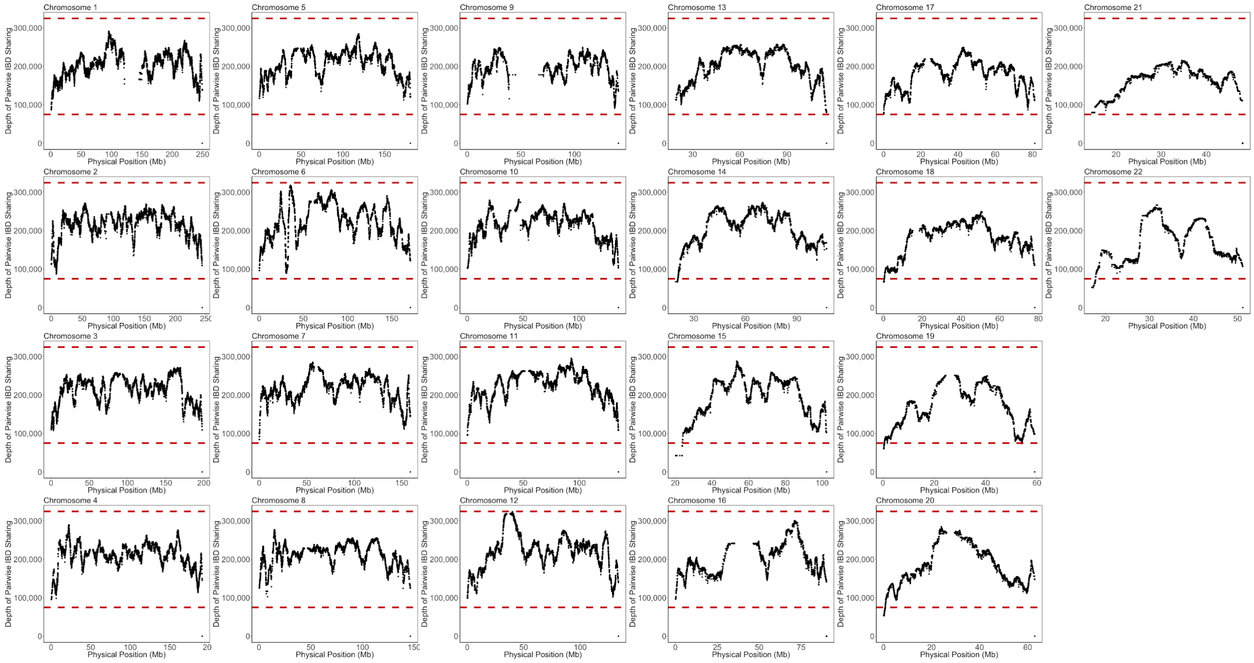

**Supplementary Figure 10. Pile up of Identity-by-Descent (IBD) haplotype sharing along the genome in Bio*Me* and TGP samples (N=34209 total). Related to Figure 3.** Each panel represents the number of shared IBD haplotypes covering a given site for each of the 22 autosomes. The x-axis represents the physical position along the chromosome (Mb) and the y-axis represents the number of haplotypes covering each position. The red dashed lines represent +/- 3 standard deviations from the genome-wide mean.

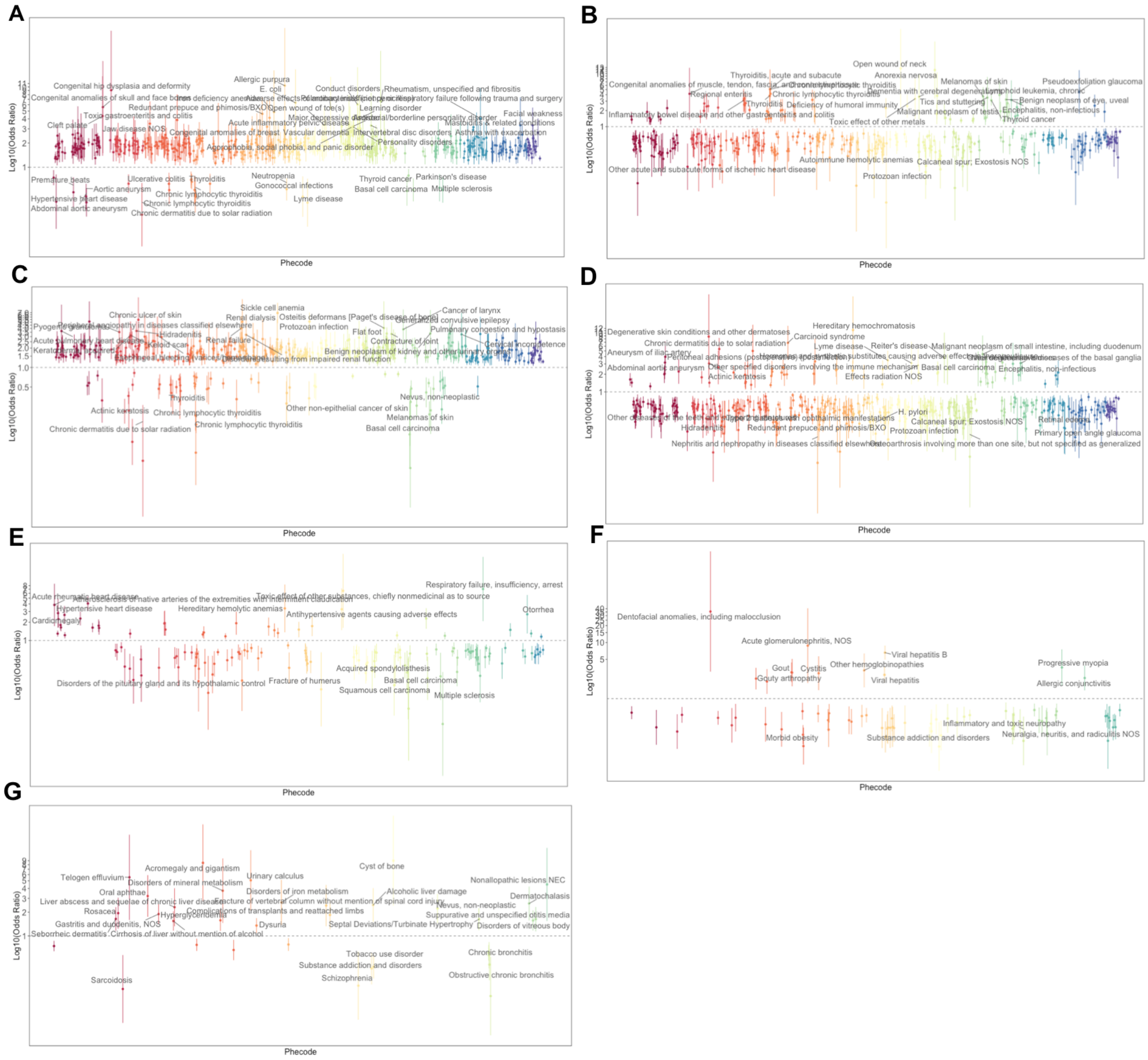

**Supplementary Figure 11. Related to Figure 5. Phenome-Wide Enrichment and Depletion of Phecodes within the 7 largest IBD Communities.** Odds ratios for the phecodes identified as being significantly enriched or depleted in the Puerto Rican (A), Ashkenazi Jewish (B), African Diaspora (C), Non-Jewish European (D), Dominican (E), Filipino (F), and Central/South American Hispanic/Latino (G) IBD-communities (odds ratios displayed are for associations that are significant after imposing a significance threshold based Bonferonni correction).

| Self-Reported Ethnicity | N | Percentage |
|---|---|---|
| African-American / African | 7976 | 22.1181 |
| American Indian / Native American | 61 | 0.169158 |
| Caucasian / White | 7961 | 22.0765 |
| East or Southeast Asian (i.e. China Japan Korea Indonesia) | 965 | 2.67602 |
| Hispanic Latin American | 11544 | 32.0124 |
| Jewish | 1869 | 5.18288 |
| Mediterranean (i.e. Spain Portugal Italy Turkey Middle East and North Africa) | 739 | 2.04931 |
| Multiple Selected | 2340 | 6.489 |
| Other | 1832 | 5.08028 |
| South Asian/Indian (i.e. India Pakistan) | 754 | 2.0909 |
| No Designation | 20 | 0.0554616 |

**Supplementary Table 1. Related to Figure 1. Self-reported ethnicity as surveyed as part of enrollment into the Bio*Me* Biobank for N=36061 participants**. Participants who selected multiple categories were grouped into the “Multiple Selected” category, with the exception of participants who self-reported as both “Jewish” and “Caucasian / White” who were grouped into the “Jewish” category. Self-reported ethnicity information was unavailable for N=20 participants.

| Ethnicity | Mean k1 | (k1 95% CI) | Mean k2 | (k2 95% CI) | Mean k3 | (k3 95% CI) | Mean k4 | (k4 95% CI) | Mean k5 | (k5 95% CI) |
|---|---|---|---|---|---|---|---|---|---|---|
| African-American / African | 0.830 | (0.827-0.833) | 0.141 | (0.138-0.144) | 0.010 | (0.009-0.011) | 0.007 | (0.007-0.008) | 0.012 | (0.011-0.013) |
| American Indian / Native American | 0.599 | (0.503-0.695) | 0.280 | (0.207-0.353) | 0.027 | (-0.009-0.064) | 0.012 | (0.005-0.019) | 0.082 | (0.023-0.141) |
| Caucasian / White | 0.015 | (0.014-0.016) | 0.921 | (0.920-0.923) | 0.041 | (0.040-0.042) | 0.006 | (0.005-0.006) | 0.017 | (0.017-0.017) |
| East or Southeast Asian (i.e. China Japan Korea Indonesia) | 0.004 | (0.002-0.006) | 0.015 | (0.011-0.020) | 0.037 | (0.028-0.045) | 0.933 | (0.921-0.944) | 0.012 | (0.011-0.013) |
| Hispanic Latin American | 0.266 | (0.262-0.271) | 0.533 | (0.529-0.537) | 0.005 | (0.005-0.006) | 0.007 | (0.007-0.008) | 0.189 | (0.185-0.193) |
| Jewish | 0.024 | (0.023-0.025) | 0.895 | (0.893-0.896) | 0.068 | (0.067-0.069) | 0.011 | (0.010-0.011) | 0.0026 | (0.001-0.004) |
| Mediterranean (i.e. Spain Portugal Italy Turkey Middle East and North Africa) | 0.044 | (0.038-0.050) | 0.866 | (0.858-0.875) | 0.077 | (0.072-0.082) | 0.004 | (0.003-0.005) | 0.0082 | (0.006-0.010) |
| Multiple Selected | 0.238 | (0.223-0.253) | 0.652 | (0.637-0.668) | 0.048 | (0.044-0.053) | 0.022 | (0.018-0.027) | 0.039 | (0.036-0.042) |
| No designation | 0.319 | (0.071-0.568) | 0.573 | (0.349-0.796) | 0.045 | (0.011-0.079) | 0.011 | (0.007-0.016) | 0.052 | (-0.040-0.143) |
| Other | 0.239 | (0.221-0.257) | 0.537 | (0.518-0.557) | 0.124 | (0.112-0.137) | 0.075 | (0.063-0.086) | 0.025 | (0.022-0.027) |
| South Asian/Indian (i.e. India Pakistan) | 0.008 | (0.004-0.013) | 0.177 | (0.166-0.188) | 0.702 | (0.689-0.715) | 0.092 | (0.081-0.104) | 0.021 | (0.020-0.021) |

**Supplementary Table 2. Related to Figure 2.** Summary of ADMIXTURE at K=5 for N=37105 Bio*Me* participants that were genotyped on the GSA. The mean and 95% confidence intervals for each k are reported, stratified by self-reported ethnicity.

| IBD Community | Size (N individuals) | Median sum IBD (cM) | IBD 95% CI | Median sum ROH (MB) | ROH 95% CI | mean k1 | mean k2 | mean k3 | mean k4 | mean k5 |
|---|---|---|---|---|---|---|---|---|---|---|
| Puerto Rican | 5452 | 18.39 | (18.38-18.41) | 10 | (9.55-10.39) | 0.26 | 0.59 | 0.00 | 0.01 | 0.14 |
| Garifuna | 113 | 121.31 | (118.28-124.95) | 41.07 | (30.60-49.75) | 0.03 | 0.89 | 0.06 | 0.01 | 0.00 |
| Colombian | 234 | 16.12 | (15.82-16.43) | 12.29 | (7.98-16.11) | 0.83 | 0.14 | 0.01 | 0.01 | 0.01 |
| Bangladeshi | 235 | 4.11 | (3.97-4.21) | 8.06 | (4.98-12.48) | 0.01 | 0.93 | 0.03 | 0.00 | 0.02 |
| Mexican | 254 | 5.56 | (5.45-5.69) | 10.04 | (7.10-13.92) | 0.40 | 0.51 | 0.01 | 0.01 | 0.08 |
| Finnish | 120 | 10.61 | (10.34-10.91) | 6.68 | (3.95-7.97) | 0.01 | 0.02 | 0.03 | 0.93 | 0.01 |
| Vietnamese | 117 | 4.17 | (4.09-4.27) | 4.71 | (3.45-5.61) | 0.09 | 0.59 | 0.01 | 0.01 | 0.30 |
| Kenyan | 101 | 8.01 | (7.76-8.25) | 4.67 | (3.80-5.77) | 0.00 | 0.01 | 0.00 | 0.96 | 0.03 |
| Ashkenazi Jewish | 4415 | 21.79 | (21.77-21.80) | 10 | (9.66-10.38) | 0.06 | 0.34 | 0.01 | 0.01 | 0.59 |
| Tuscani | 102 | 6.06 | (5.75-6.39) | 5.46 | (4.11-9.49) | 0.80 | 0.05 | 0.00 | 0.00 | 0.15 |
| African Diaspora | 7470 | 4.62 | (4.61-4.63) | 4.02 | (3.91-4.21) | 0.09 | 0.60 | 0.01 | 0.01 | 0.30 |
| Non-Jewish European | 6183 | 4.06 | (4.06-4.07) | 4.27 | (4.09-4.44) | 0.00 | 0.09 | 0.74 | 0.15 | 0.02 |
| Dominican | 1971 | 11.81 | (11.78-11.84) | 13.52 | (12.02-15.05) | 0.03 | 0.16 | 0.01 | 0.02 | 0.78 |
| Filipino | 699 | 4.25 | (4.22-4.30) | 5.86 | (4.53-6.86) | 0.01 | 0.88 | 0.03 | 0.04 | 0.04 |
| Central/South American | 1265 | 4.44 | (4.42-4.47) | 6.37 | (5.23-6.81) | 0.00 | 0.00 | 0.03 | 0.96 | 0.00 |
| Korean/Japanese | 364 | 3.69 | (3.66-3.72) | 3.91 | (3.60-4.52) | 0.34 | 0.56 | 0.09 | 0.01 | 0.00 |
| Ecuadorian | 438 | 7.69 | (7.57-7.81) | 8.77 | (6.98-11.53) | 0.02 | 0.93 | 0.04 | 0.01 | 0.01 |

**Supplementary Table 3. Related to Figure 4. Summary Statistics for the 17 largest communities (N >100) detected via *InfoMap*.** The first column represents how many participants fall into each IBD-community, followed by the distribution of the sum of Identity-by-Descent (IBD) sharing (community-wide median and 95% confidence intervals), along with the abundance of Runs of Homozygosity (ROH) (MB) (median and 95% C.I.). The final 5 columns represent the mean global continental ancestry components per community, as estimated *via* ADMIXTURE at k=5.

| **Community** | **Clustering Coefficient** | **Wasserstein Metric** |
|---|---|---|
| Puerto Rican | 0.852 | 0.780 |
| Garifuna | 0.972 | 0.960 |
| Colombian | 0.789 | 0.690 |
| Bangladeshi | 0.084 | 0.035 |
| Mexican | 0.301 | 0.201 |
| Finnish | 0.832 | 0.791 |
| Vietnamese | 0.169 | 0.148 |
| Kenyan | 0.732 | 0.696 |
| Ashkenazi Jewish | 0.918 | 0.821 |
| Tuscani | 0.340 | 0.235 |
| African Diaspora | 0.053 | 0.032 |
| Non-Jewish European | 0.090 | 0.027 |
| Dominican | 0.594 | 0.449 |
| Filipino | 0.215 | 0.054 |
| Central/South American | 0.088 | 0.027 |
| Korean/Japanese | 0.139 | 0.063 |
| Ecuadorian | 0.380 | 0.221 |

**Supplementary Table 4. Related to Figure 4. Clustering coefficient and Wasserstein metric for the 17 largest communities (N >100) detected via *InfoMap*.** The first column lists the clustering coefficient, defined as the ratio of existing edges between the neighbors of a specific node relative to the total number of potential edges between said neighbors. The second column lists the Wasserstein metric computed between the intra-community degree distribution and the inter-community distribution.

**Supplementary Table 5. Related to Figure 5.** Table representing all statistical associations between IBD-community membership for the seven largest IBD-communities and phecodes that passed Bonferroni correction. **[***Supplementary Table 5 was uploaded as a separate spreadsheet***].**

| ***"Mark each of the ethnic groups that are part of your family background. You may select more than one:"*** |
|---|
| *"Hispanic Latin American"* |
| *"African-American / African"* |
| *"Caucasian / White"* |
| *"Jewish"* |
| *"Mediterranean (i.e. Spain Portugal Italy Turkey Middle East and North Africa)"* |
| *"American Indian / Native American"* |
| *"East or Southeast Asian (i.e. China Japan Korea Indonesia)"* |
| *"South Asian/Indian (i.e. India Pakistan)"* |
| *"Other"* |

**Supplementary Table 6. Related to Figure 1. Survey of self-reported ethnicity as represented in the Bio*Me* biobank questionnaire.** This question is administered to Bio*Me* participants at enrollment as part of a larger questionnaire. The question is presented as multiple choice, allowing for participants to select more than one category.
